# In situ molecular architecture of PML bodies reveals open-state columnar trinucleosome assemblies within a porous, chromatin-permissive interior

**DOI:** 10.64898/2026.06.07.729032

**Authors:** Vojtěch Pražák, Iain Harley, Julian Falckenhayn, Chris Boutell, Peter A. Thomason, Benjamin G. Davis, Rainer Kaufmann, Stephen D. Carter

## Abstract

Genome function in the nucleus is organised through membrane-less compartments enriched with specific proteins and nucleic acids. Promyelocytic leukemia (PML) bodies regulate telomere maintenance and DNA damage responses, but their internal molecular organisation remains poorly understood. We visualised the molecular composition of PML bodies directly within the nucleus using an advanced cryogenic-correlative light and electron microscopy (cryo-CLEM) pipeline. Cryo-electron tomography revealed eYFP-PML-I bodies as compartments with a molecular makeup distinct from the surrounding nucleoplasm. Cryogenic super-resolution correlative light and electron microscopy showed these comprise a diffuse PML-protein shell enclosing an inner core. A visual proteomics comparison of the core and surrounding regions through template matching and sub-tomogram averaging mapped nucleosomes, TRiC chaperonin complexes in both closed and open conformations, and single-capped PA28 proteasomes. Membrane structures of unknown function were additionally detected within some eYFP-PML-I bodies. Remarkably, the nucleosomes included density consistent with columnar trinucleosome assemblies (1), revealing that PML body interiors harbour discrete chromatin domains. Moreover, vault complexes were detected in poised positions in the proximal nucleoplasm. Together, these first in situ higher-resolution structural insights of PML bodies identify their critical functional components and an interior porous architecture that suggests a mechanism for content selection based on physical partitioning akin to size-exclusion chromatography.

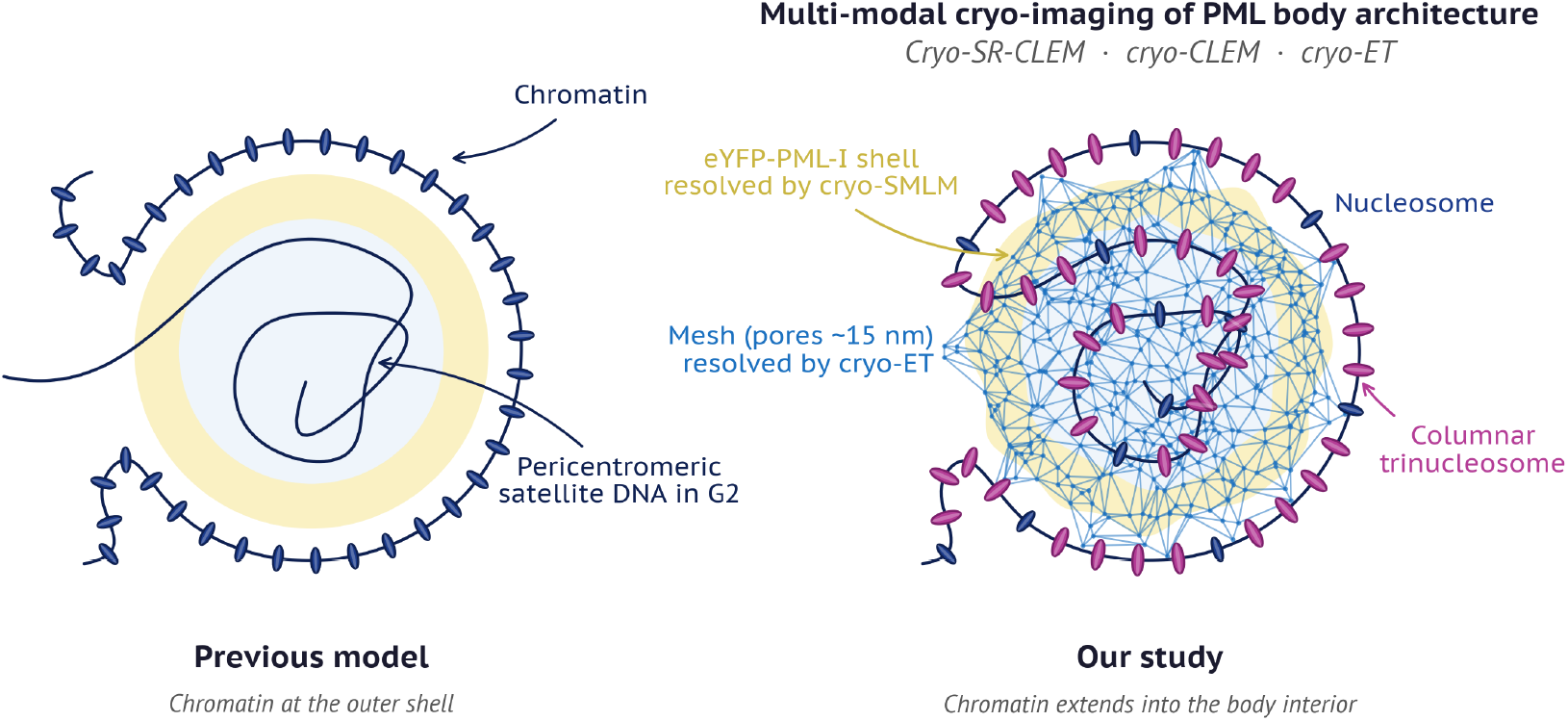

## Introduction

The eukaryotic nucleus achieves its spatial organisation through selective separation, generating membrane-less condensates. Some of these, including nucleoli, nuclear speckles, paraspeckles and classical Cajal bodies (2, 3), assemble through multivalent protein-protein and protein-RNA interactions, often involving the intrinsically disordered regions (IDRs) of RNA-binding proteins. Insights into the resulting specialised biochemical environments have given insight into the way that they regulate nuclear processes (4). By contrast, other important nuclear bodies are much less characterised; foremost amongst these are the promyelocytic leukemia (PML) bodies that assemble through exploitation of SUMO (small ubiquitin-like modifier)–SIM (SUMO-interacting motif) networks, forming major hubs that can concentrate over 200 client proteins (5, 6). Functionally, PML bodies orchestrate telomere maintenance, cellular senescence, protein degradation, the DNA damage response, gene transcription, apoptosis and antiviral defence (7, 8) yet the underlying mechanisms and, indeed, the structure that accomplishes this remains opaque.

PML bodies were first observed by conventional electron microscopy in the 1960s as electron-dense interchromatin nuclear bodies (9, 10). These structures gained molecular identity when cloning of PML from the acute promyelocytic leukemia (APL) translocation (11, 12) was followed by immuno-labelling electron microscopy studies demonstrating that PML protein localized specifically to these nuclear bodies (13, 14). The observation that these bodies were disrupted in APL and reformed upon treatment has provided an archetypal example of targeted cancer therapy (15).

The characteristic PML body architecture comprises a ∼100 nm PML-containing shell surrounding a PML-free inner core (16, 17). Super-resolution microscopy, particularly 4Pi fluorescence imaging (18), suggests that PML and Sp100 proteins form distinct, partially overlapping patches within this shell, potentially enabling dynamic protein exchange with the nucleoplasm (19). This ring-like shell architecture has also been shown by 3D-SIM (20). One central PML protein, TRIM19, belongs to the tripartite motif (TRIM) family characterised by an N-terminal RING domain, two B-box zinc finger domains, and a coiled-coil region. This architecture resembles that of TRIM5α, which restricts retroviral infection and contains only a single B-box domain (21).

While PML body cores are generally considered protein-rich structures, DNA incorporation into specialised subtypes has also been reported. Alternative lengthening of telomeres (ALT)- associated PML bodies (APBs) in telomerase-negative cancer cells contain telomeric DNA that supports recombination-based telomere maintenance (22, 23), while satellite DNA accumulates in PML bodies during G2 to promote heterochromatin re-modelling (24). Giant PML bodies in ICF (immunodeficiency, centromeric instability and facial anomalies) syndrome contain under-condensed satellite DNA cores (25), and latent herpesvirus genomes are sequestered within viral PML bodies as a seeming antiviral mechanism (26, 27). Together, these observations suggest that DNA content of PML bodies may be dynamic and functionally regulated. Notably, although PML bodies have been suggested to associate with chromatin (17, 19, 20, 28), direct evidence such as through visualisation of nucleosomes within their cores has not been obtained. Conventional imaging approaches lack the resolution to determine the precise organisation of DNA within these condensates. Moreover, any higher-order chromatin architecture of possible telomeric DNA associated with PML bodies remains equally unresolved. More broadly, the extent to which macromolecular complexes are selectively partitioned into or excluded from PML body interiors has not been examined at the structural level, leaving a gap in our understanding of the mechanism that underpins their dynamic roles within the nucleus as regulatory hubs.

Here, we show that a cryogenic correlative light and electron microscopy (cryo-CLEM) pipeline using plasma focused ion beam (pFIB) milling can visualise PML bodies at molecular resolution within vitrified nuclei. Using cryo-electron tomography (cryo-ET), template matching and sub-tomogram averaging (STA), this now reveals through direct visualisation, DNA in nucleosomes including open-state columnar trinucleosome assemblies. This previously unknown chromatin organisation within PML body interiors demonstrates selective partitioning of macro-molecular complexes and suggests that this is achieved via a physical mechanism involving a porous interior mesh akin to size-exclusion chromatography.

## Results

### Cryo-CLEM and associated textural differences enable efficient identification of PML bodies within nuclei

To characterise the molecular architecture of PML bodies, we employed human foreskin fibroblasts (HFFs) stably expressing doxycycline-inducible eYFP-PML-I (hereafter PML-I) (29, 30). HFFs exhibited exceptional thin morphology when cultured on EM grids, making them ideal for accessing deep nuclear regions where PML bodies reside. Cells were seeded on gold SiO_2_ film grids for 12 hours. Following a 7-hour doxycycline induction, sufficient for eYFP-PML-I to incorporate into existing endogenous PML bodies (Fig. 1A), cells with bright nuclear foci were identified by an integrated fluorescence light microscope (iFLM) and their positions registered in x, y coordinates using an integrated scanning electron microscope (SEM).

**Figure 1.**
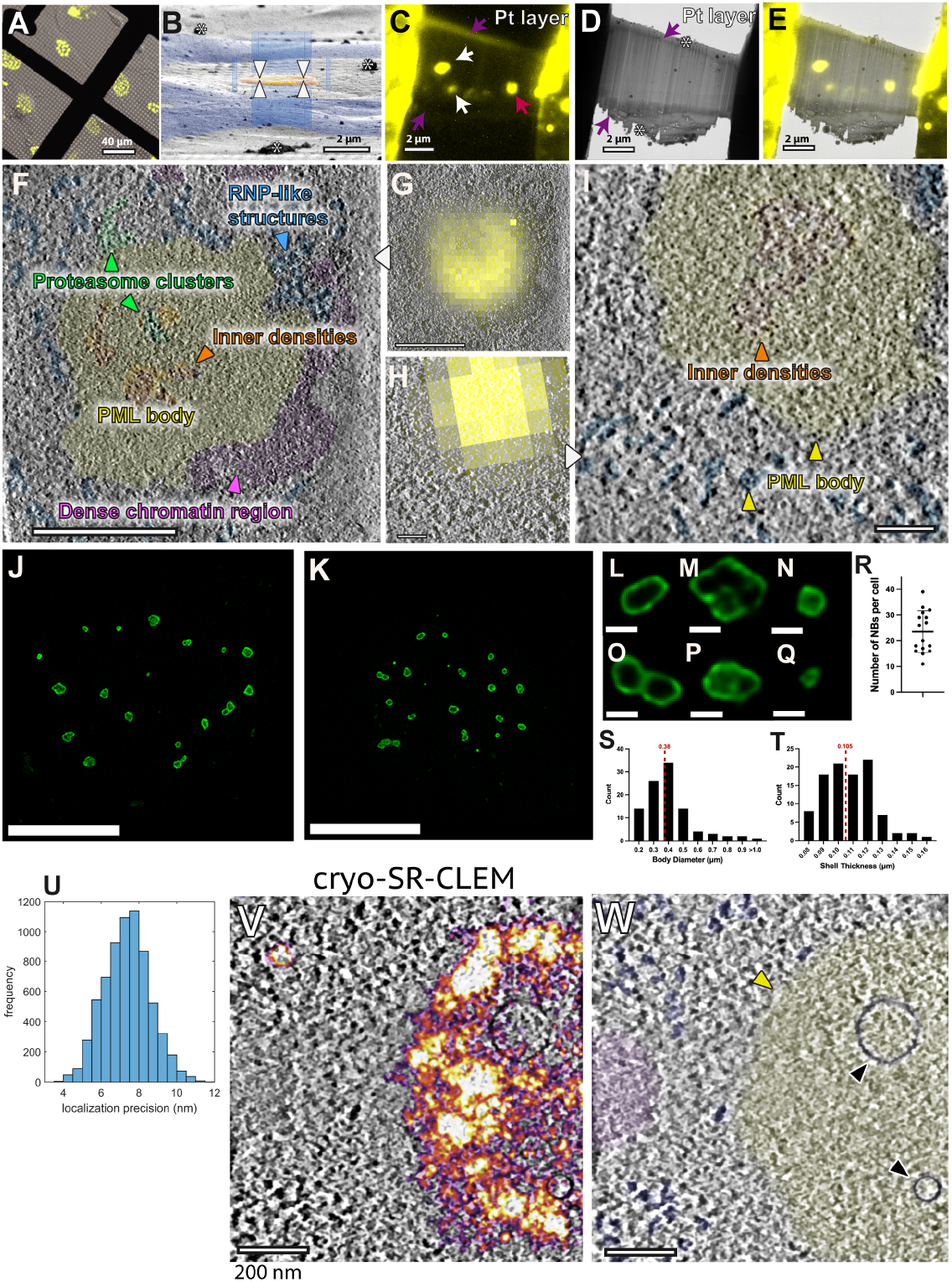
Cryo-CLEM pipeline for in situ visualisation of eYFP-PML-I bodies. **(A)** A representative cryo-fluorescence image of human foreskin fibroblasts (HFFs) stably expressing doxycycline-inducible eYFP-PML-I cultured on gold SiO2 film grids. eYFP-PML-I bodies appear as yellow puncta. Scale bar = 40 µm. **(B)** Plasma ion image of vitrified HFFs. The nucleus is shaded in orange. White arrowheads indicate lamella targeting position. Scale bar = 2 µm. **(C)** Post-milling cryo-fluorescence image of the lamellae. Purple arrows indicate autofluorescence of the platinum (Pt) deposition layer, used as landmarks for cryo-CLEM. White arrows indicate other eYFP-PML-I bodies captured within the lamellae. Pink arrow indicates the eYFP-PML-I body shown in (F and G). Scale bar = 2 µm. **(D)** Cryo-EM search map of the same lamellae. Purple arrows indicate the platinum layers. Scale bar = 2 µm. **(E)** Overlay of cryo-fluorescence and cryo-EM search map, enabling precise correlation of eYFP-PML-I signal for tilt-series collection. Scale bar = 2 µm. **(F)** Denoised cryo-tomogram slice of the eYFP-PML-I body collected at 33,000× magnification indicated by the pink arrow in **(C).** Green arrowheads indicate proteasome clusters; blue arrowhead indicates ribonucleoprotein (RNP)-like structures; orange arrowhead indicates inner densities; pink arrowhead indicates a dense chromatin region. Scale bar = 500 nm. **(G)** Cryo-fluorescence of the eYFP-PML-I body from **(C)** overlaid onto the cryo-tomogram slice, showing correlation of fluorescence signal with the finely textured region. Scale bar = 500 nm. **(H)** A different eYFP-PML-I body collected at higher magnification (53,000×) with cryo-fluorescence overlaid onto the cryo-tomogram slice. Scale bar = 150 nm. **(I)** Cryo-tomogram slice through the eYFP-PML-I body in H. Orange arrowhead indicates inner densities; yellow arrowhead indicates the PML body; blue arrowhead indicates RNP-like structures. Scale bar = 150 nm. **(J–K)** 3D-SIM images of HFFs expressing eYFP-PML-I, showing characteristic hollow shell architecture of PML bodies across the nucleus. Scale bars = 10 µm. **(L–Q)** Zoomed 3D-SIM views of individual eYFP-PML-I bodies illustrating morphological diversity: **(L)** classical round body; **(M)** large body with multiple core density chambers; **(N)** small round body; **(O)** figure-of-eight morphology; **(P)** round body with internal density; **(Q)** small body with no discernible core. Scale bars = 500 nm. **(R)** Scatter plot showing the number of PML nuclear bodies per cell (n = 16 cells). **(S)** Histogram of eYFP-PML-I body diameters measured from 3D-SIM data. Median = 380 nm (red dashed line). **(T)** Histogram of eYFP-PML-I shell thickness measured from 3D-SIM data. Median = 105 nm (red dashed line). **(U)** Histogram of single-molecule localisation precision from cryo-SR-CLEM data. **(V)** cryo-SR-CLEM localisations overlaid onto a cryo-ET tomogram slice, demonstrating that the eYFP-PML-I shell resides inside the finely textured region that lacks large granular densities. Scale bar = 100 nm. **(W)** Cryo-tomogram slice with annotation. Yellow arrowhead indicates the PML body; black arrowheads indicate membrane vesicles. Scale bars = 200 nm.

Next, we established a cryo-CLEM workflow in which the eYFP-PML-I fluorescence signal was used to guide lamella positioning (Fig. 1B), followed by on-lamella correlation to target tilt-series acquisition (Fig. 1C-E). Lamellae were generated within fluorescent nuclei (Fig. 1B); after milling, approximately 40% of lamellae retained their detectable fluorescence (Fig. 1C), enabling reliable correlation between cryo-fluorescence and electron microscopy images for precise targeting (Fig. 1D-E). Using this approach, we collected tilt-series both directly on PML-I bodies and in adjacent nuclear regions to representatively sample the surrounding nucleoplasm. In total, from 13 vitreous lamellae we acquired 45 tilt-series targeting 27 PML bodies and 18 neighbouring regions, from which 7 PML body cryo-tomograms and 9 nucleoplasmic cryo-tomograms were selected for high-resolution analysis (Sup. Fig. 5A and B).

The visualisation of membrane-less compartments within cells including inside the nucleus remains comparatively challenging, even in high-resolution cryo-ET data, as they lack clear boundaries and appear featureless relative to e.g. the surrounding nucleoplasm. Advantageously, in our cryo-tomograms, it proved possible to identify PML-I bodies due to an identifiable difference in texture compared to the surrounding nucleoplasm. Assignment of these 400-800 nm diameter structures was provided by the cryo-iFLM signal initially used for targeting regions for high-resolution data acquisition (Fig. 1E, G and H, 2F). Further use of IsoNet denoising (31) and total variation denoising (32) revealed a clear, irregularly shaped boundary between the edge of putative PML bodies and granular nucleoplasm (Fig. 1F and I, 2G, 3E). The interiors contained a fine-grained texture distinct from the larger granular appearance of the surrounding nucleoplasm (Fig. 1F and I), reminiscent of TRIM5α-bodies observed by previous cryo-ET work (33) (see Discussion).

### 3D-SIM and cryo-SR-CLEM confirm PML-I body shell architecture

To characterise PML-I body dimensions and morphology under the same culture conditions used for cryo-ET, we performed 3D-SIM on the same HFF cell line. 3D-SIM confirmed the characteristic hollow shell architecture of PML-I bodies (20) (Fig. 1J, K), but also considerable morphological diversity including classical round bodies, and previously unknown multi-chambered bodies, figure-of-eight configurations, and bodies with internal densities (Fig. 1L–Q). Quantitative analysis across 16 cells gave a median diameter of 380 nm and a median outer shell thickness of 105 nm (Fig. 1R–T), consistent with the finely textured regions observed in our cryo-tomograms containing PML-I bodies. To precisely colocalise the PML-I shell within these cryo-ET structures, we performed cryogenic super-resolution correlative light and electron microscopy (cryo-SR-CLEM) (34), observing single eYFP-PML-I molecules with an average precision of 7.7 nm (Fig. 1U). Fluorescence signal concentrated at the periphery of these finely textured regions (Fig. 1V–W), thereby placing the eYFP-PML-I shell at their outermost boundary. Interestingly, while 3D-SIM suggests a smooth PML-I protein boundary, cryo-SR-CLEM reveals this to be irregularly contoured at the nanoscale, consistent also with the irregular boundary visible in cryo-ET between the finely textured body region and the surrounding nucleoplasm. The PML-I protein shell itself, however, is not resolved as a distinct layer in cryo-ET, consistent with the shell being assembled from many small proteins whose densities blend continuously into both the body interior and the surrounding granular nucleoplasm.

### PML-I body interiors are compositionally and texturally distinct from the surrounding nucleoplasm

To compare the molecular composition of PML-I body interiors with the surrounding nucleoplasm, we first delineated the body boundaries within each cryo-tomogram. We segmented PML-I body regions using cryo-CLEM data and denoised tomograms with manual segmentation in Amira (35) to define the boundary between PML-I bodies and the surrounding granular nucleoplasm (Fig. 2T).

**Figure 2.**
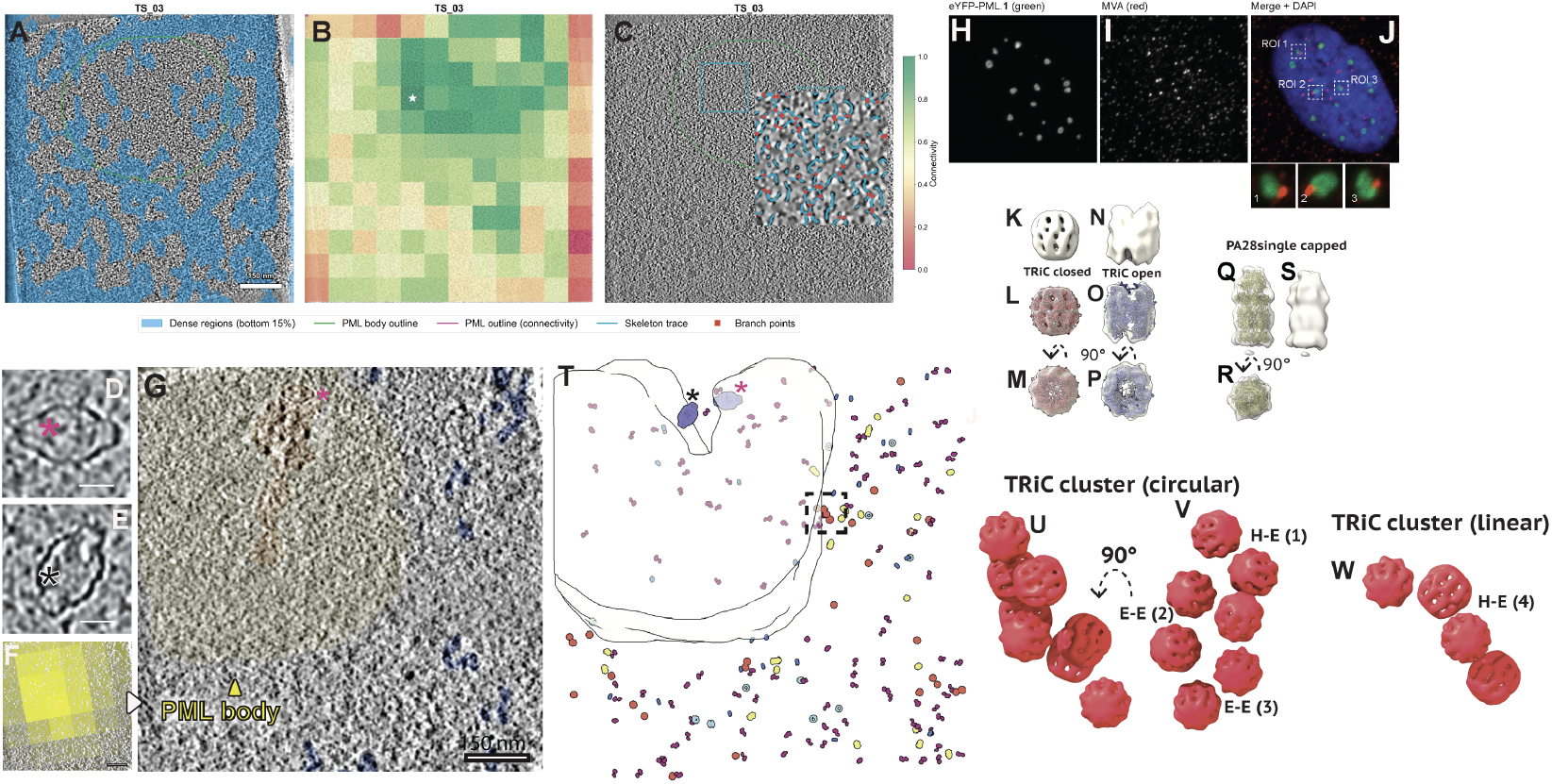
Interior network analysis and visual proteomics of macromolecular complexes in eYFP-PML-I cryo-tomograms. **(A)** Z-projection of TS_03 showing high-contrast dense regions (cyan, bottom 15% intensity) relative to the segmented PML body (green outline). Dense regions are spatially excluded from the PML body interior. **(B)** Skeleton connectivity heatmap (12×12 grid) overlaid on mean-intensity projection of TS_03. Connectivity score (0–1, red–green) was computed from black top-hat segmentation followed by 2D skeletonization of each slice, combining fibre density (60%) and branch point density (40%). White star indicates the most connected grid cell. **(C)** Representative cryo-tomogram slice of TS_03 with skeleton trace inset. Cyan box indicates the most connected grid region; inset shows the denoised crop with skeleton overlay (cyan) and branch points (red). **(D)** Cryo-tomogram slice showing a vault complex indicated by a pink asterisk in G and T. Scale bar = 25 nm. **(E)** Cryo-tomogram slice showing the vault complex indicated by the black asterisk in **(T).**Scale bar = 25 nm. **(F)** Cryo-fluorescence of the eYFP-PML-I body overlaid onto the cryo-tomogram slice. Scale bar = 150 nm. **(G)** Cryo-tomogram slice of the eYFP-PML-I body showing the PML body (yellow arrowhead) and the vault complex (pink asterisk). Scale bar = 150 nm. **(H)** eYFP-PML-I channel (green). **(I)** MVP immunofluorescence (red). **(J)** Merged image with DAPI showing colocalisation of eYFP-PML-I and MVP signals across ROIs 1–3 (dashed boxes), with zoomed insets. **(K)** STA map of TRiC in the closed conformation. **(L)** Same as **(K)** rendered transparent with PDB 7 × 6Q fitted. **(M)** Same as **(L)** rotated 90°. **(N)** STA map of TRiC in the open conformation. **(O)** Same as **(N)** rendered transparent with PDB 7 × 7Y fitted. **(P)** Same as **(O)** rotated 90°. **(Q)** STA map of PA28 single-capped proteasome rendered transparent with atomic model fitted (PDB 7DR6). **(R)** Same as **(Q)** rotated 90°. **(S)** Solid surface map of the PA28 single-capped proteasome. **(T)** STA particles mapped back onto TS_01. Closed TRiC (red), open TRiC (light blue), and PA28 single-capped proteasomes (yellow) are mapped onto the segmented PML body mesh (white). Trinucleosomes are shown in magenta; nucleosomes in blue. Pink and black asterisks indicate the vault complexes shown in **(D)** and **(E)**, respectively. Dashed box indicates linear TRiC cluster shown in W. **(U)** TRiC cluster in a circular arrangement, shown without labels. **(V)** Same cluster as **(U)** rotated 90°, with labelled contact geometries: head-to-equator (H-E, type 1) and equator-to-equator (E-E, types 2 and 3; see also Sup. Fig. 8H). **(W)** TRiC cluster in a linear arrangement (H-E, type 4).

To assess the distribution of large macromolecular densities, we identified high-contrast dense regions and compared their abundance inside and outside the PML-I body (see Methods; Fig. 2A, Sup. Fig. 2A). High-contrast dense material occupied only 0.7 to 3.2 percent of the PML-I body interior, compared with 11 to 12 percent of the surrounding nucleoplasm across the three cryo-tomograms analysed, a 3- to 17-fold depletion (4.2-fold in TS_01, 17.1-fold in TS_03 and 3.4-fold in TS_08) indicating spatial exclusion of large granular densities from PML-I body interiors. Skeletonisation (36) of the internal density network further revealed consistently higher connectivity within PML body regions compared to the surrounding nucleoplasm in all three cryo-tomograms (Fig. 2B, C, Sup. Fig. 2B–D), indicating that PML body interiors maintain a denser, more extensively branched mesh-like network than the adjacent nucleoplasm. Direct measurement of gap sizes in the skeletonised interior network gives a mean pore diameter of 15.4 nm (standard deviation 5.7 nm) across 363,437 pooled pores from the three cryo-tomograms analysed (Sup. Fig. S3).

### Molecular architecture within PML-I cryo-tomograms

Two cryo-tomograms revealed intact vault complexes adjacent to the periphery of the finely textured regions (Fig. 2D, E, G and T). Immunofluorescence using an antibody against the major vault protein (MVP) revealed discrete puncta adjacent to PML-I bodies in multiple cells (Fig. 2H–J), consistent with these rare events reflecting a genuine spatial association of vault complexes near PML-I bodies. As a positive control, immunofluorescence against Daxx, a known PML body component, showed clear colocalisation with PML-I bodies (Sup. Fig. S4A–C), confirming the validity of our MVP immunofluorescence approach. We also observed membrane vesicles within (n=2) and adjacent to (n=2) PML-I bodies (Fig. 1V, W).

Not only did we observe large complexes such as vaults, but we also identified a wide variety of other densities including capsules, granules, filaments, cylindrical structures, large barrels, and rings (Sup. Fig. S1C–H), as well as densities consistent with nucleosomes. To identify and characterise these macromolecular complexes, we performed template matching on 16 cryo-tomograms with thickness between 134 nm and 230 nm. Of these, 7 cryotomograms captured eYFP-PML-I fluorescence directly, while 9 captured regions adjacent to PML-I bodies at eYFP-PML-I centroid to cryo-tomogram centroid distances ranging from 780 nm to 4 μm (Sup. Fig. 5B). We performed template matching using PyTOM v0.7.1 on 4× binned tomograms (pixel size 9.92 Å). Templates were derived from published cryo-EM density maps or atomic models (EMDB/PDB; Sup. Fig. 6). Both templates and cryo-tomograms were low-pass filtered to 40 Å prior to template matching to minimise template bias. Correlation score maps were inspected and a cross-correlation threshold selected using a custom napari (37) viewer (see Methods), and candidate particles above this threshold were identified. Where multiple templates were applied to a single complex type, overlapping detections within 10 nm were resolved by cross-correlation (CC) score competition, assigning each spatial position to the highest-scoring template. Selected particle sets were imported into RELION 4.0 for STA, with iterative 3D classification applied to each dataset. Template sources, initial detection counts, particles imported into RELION 4.0, final particle numbers after 3D classification, and STA resolutions for all complexes are summarised in Sup. Fig. 6.

### TRiC chaperonin complexes form conformation-dependent clusters in the nucleus

Visual inspection of the cryo-tomograms revealed capsule-shaped densities approximately 17 nm in diameter, sometimes forming clusters (Sup. Fig. 1C). This morphology and size are consistent with TRiC/CCT chaperonin complexes, which form a characteristic double-ring barrel approximately 17 nm in diameter. We therefore searched for TRiC using two templates derived from published structures (EMDB-18913, closed state; EMDB-18921, open state). Template matching against all 16 tomograms yielded 378 closed-state and 580 open-state particle candidates (Sup. Fig. 7A). Since both templates have similar features and dimensions, many positions were detected by both. CC score competition resolved 210 overlapping positions, with closed TRiC winning 167 of these (79.5%), consistent with its higher mean CC score (0.472 vs 0.398). This yielded final particle sets of 335 closed and 413 open TRiC particles (Sup. Fig. 7B). STA in RELION 4.0 produced final maps of 297 closed and 276 open TRiC particles at 22 Å and 32 Å resolution respectively (Fig. 2K–P, Sup. Fig. 7C-F). When mapped back onto the cryo-tomograms, the closed- and open-state particles corresponded to densities consistent with closed and open TRiC, respectively (Sup. Fig. 8A–D). The double-ring barrel architecture was clearly resolved in both maps, with the central cavity visible. The closed map showed a compact, symmetric arrangement (Fig. 2K–M) while the open map displayed characteristic splayed apical domains (Fig. 2N–P).

To investigate the spatial organisation of TRiC in situ, we performed Density-Based Spatial Clustering of Applications with Noise (DBSCAN) (38) on the 3D coordinates of each conformational class (eps ≈ 37 nm; min_samples = 2). This eps value was chosen to capture extended clusters where particles are potentially linked through chains of neighbours. Closed TRiC exhibited a striking propensity for self-association, with 29.0% of particles (88/303) assigned to 35 distinct clusters (Fig. 2T; Sup. Fig. 8E, F; Supplementary Movie 1). In marked contrast, only 5.8% of open TRiC particles (16/276) formed clusters, yielding just 8 clusters (Sup. Fig. 8F). This five-fold difference in clustering frequency (Fisher’s exact test, p = 6 × 10□^14^, risk ratio 5.0, 95% CI 3.0–8.3) revealed that TRiC self-association is strongly conformation-dependent and agrees with previous cryo-ET reports of TRiC clustering in the cytoplasm (39). In that context, closed-state clustering was shown to be substrate-dependent, diminishing upon translation inhibition, suggesting that conformation-dependent clustering reflects the spatial co-organisation of active protein folding (39).

Cluster size analysis revealed that the majority of closed TRiC clusters were dimers (27/35, 77%), with additional trimers (n = 3), tetramers (n = 3), a pentamer (n = 1), and an octamer (n = 1) (Sup. Fig. 8F). Open TRiC clusters were exclusively dimeric (8/8, 100%). To characterise the packing geometry of TRiC clusters, we measured centre-to-centre distances between mutual nearest-neighbour pairs identified using a 20 nm distance cutoff (39). For closed TRiC, the mean nearest-neighbour distance was 17.3 nm (median 17.4 nm, s.d. = 0.9 nm, n = 32 pairs), closely matching the TRiC particle diameter of ∼17 nm (Sup. Fig. 8G). The two open TRiC pairs identified at this distance threshold were separated by ∼16.7 nm, although with only two pairs this open-state geometry is anecdotal (Sup. Fig. 8G).

We next examined the relative orientations of paired TRiC complexes using their Euler angles to determine the barrel axis of each particle (see Methods). Closed TRiC angles were taken from D8-symmetry-refined STA, while open TRiC angles were from C1 refinement. Of the 32 closed TRiC pairs within 20 nm, 10 (31%) adopted an equator-to-equator arrangement with aligned barrel axes and 22 (69%) a head-to-equator arrangement, with no true head-to-head contacts. Most of these head-to-equator pairs had near-perpendicular barrel axes, while five of the 22 were aligned-barrel configurations assigned to an intermediate class (Sup. Fig. 8H). Both open TRiC pairs were likewise head-to-equator (Sup. Fig. 8H).

Visual inspection of representative clusters in our cryo-tomograms confirmed these contact geometries, also revealing both circular (Fig. 2U, V) and linear (Fig. 2W) TRiC arrangements. We next mapped TRiC particle positions onto the segmented PML-I body meshes (Fig. 2T; see Methods) to determine whether complexes were localised within the body interior or restricted to the surrounding nucleoplasm. Whilst variable across tomograms (2.6% in TS_01 to 38.3% in TS_15), few TRiC complexes (29 of 194, 14.9%) were observed within PML-I body interiors (Sup. Fig. 9 and 10), with the representative example in Fig. 2T containing one TRiC complex within the segmented body region. These findings indicate that PML-I body interiors are not the predominant sites of TRiC-mediated protein folding. However, to determine whether proximity to PML bodies influences TRiC activity in the surrounding nucleoplasm, we compared closed TRiC clustering across all 16 tomograms, stratified by whether each dataset contained an eYFP-PML-I-captured body or was collected from distal nucleoplasm (Sup. Fig. 5B). Closed TRiC complexes showed a trend towards more frequent local spatial clustering (DBSCAN, eps ≈ 37 nm, min_samples = 2) in PML-body-proximal tomograms (mean 36.6% of particles per tomogram, range 17.4–66.7%; clusters detected in all 7 datasets) than in distal nucleoplasmic tomograms (mean 17.1%, range 0–55.6%; clusters detected in 5 of 9 datasets), although this difference did not reach statistical significance (Mann-Whitney U test, two-sided p = 0.07). This is consistent with, but does not establish, a higher frequency of closed-TRiC association near PML-I bodies, and whether it reflects enhanced protein folding activity remains to be determined.

### Identification of PA28 single-capped proteasomes by multi-template matching

Our cryo-tomograms also revealed cylindrical densities approximately 15 nm in diameter, consistent with the barrel-shaped 20S proteasome core particle. Since PA28-capped proteasomes have previously been suggested to accumulate at PML bodies (40), we hypothesised these could represent proteasome complexes. We applied template matching in PyTOM using seven proteasome-related templates including single- and double-capped variants of PA28, PA200, and 19S, as well as the uncapped 20S core (Sup. Fig. 6; Supplementary Note 1). CC score analysis and spatial overlap grouping (Supplementary Note 1) indicated that double-capped and 19S species were not detected across any of the 16 cryo-tomograms.

Single-capped PA28 consistently achieved the highest CC scores of all cap-containing templates (Supplementary Note 1; Sup. Fig. 6); however, cap-alone detections are susceptible to false positives from other cellular densities of similar size, necessitating a two-template spatial validation strategy. To recover genuine PA28 proteasomes, we therefore required each cap detection to spatially overlap with a 20S Core detection within 10 nm (Sup. Fig. 11). This two-template filter yielded 875 validated PA28 single-capped proteasome particles. STA in RELION 4.0 with iterative 3D classification yielded a final reconstruction of 389 particles at 29 Å resolution (Fig. 2Q–S; Sup. Fig. 12C–F), with clear densities for both the 20S core barrel and the PA28 cap when particles were plotted back onto cryo-tomograms (Sup. Fig. 12D–E). We also tested an alternative pipeline incorporating PA28/PA200 CC competition and orientation filtering (Phase 2), which yielded fewer PA28-winner particles at lower resolution (33 Å; Sup. Fig. 12A–B). Independent STA of PA200-winner particles produced uninterpretable density consistent with false-positive cross-matching, and PA200 was not pursued further.

To determine the spatial distribution of PA28 single-capped proteasomes relative to PML-I body architecture, we mapped final PA28 particle positions onto the five segmented PML body meshes (Sup. Fig. 9 and 10). The majority of PA28 particles were localised outside the PML body interior across all five segmented tomograms: TS_01 (0%, 0/18), TS_03 (11.5%, 3/26), TS_08 (3.4%, 3/88), TS_15 (25.9%, 7/27), and TS_16 (0%, 0/8). Overall, only 13 of 167 PA28 particles (7.8%) were located within PML bodies, indicating that PA28 single-capped proteasomes are excluded from the PML body interior more strongly than TRiC (14.9% inside). DBSCAN analysis of 389 PA28 particles across all 16 cryo-tomograms revealed limited spatial clustering, with the two largest clusters comprising 52 and 15 particles (Sup. Fig. 13A-C).

### Molecular architecture of nucleosome core particles within PML-I cryo-tomograms

Having established that our template matching and STA pipeline was effective for relatively large macromolecular complexes, we next searched for nucleosome core particles using a template derived from cryo-EM (EMDB-8140), to determine whether chromatin occupies PML-I body interiors, a question that has remained unresolved at molecular resolution.

STA of 5,517 particles in RELION 4.0, through iterative alignment and extensive 3D classification, yielded a reconstruction of the nucleosome core particle (Fig. 3A-C) at 19 Å resolution after two rounds of 3D classification (957 particles; Sup. Fig. 14D, G). Notably, DNA strands were clearly visible, with sufficient detail to resolve major grooves allowing localisation of chromatin in PML bodies for the first time (Fig. 3E-F). Our 3D classification analysis also revealed 777 particles displaying an extra density adjacent to the nucleosome (Sup. Fig. 14E, F, marked with an asterisk), consistent with bound accessory proteins (41). When particle positions were mapped back onto the cryo-tomograms, both classes aligned well with high-contrast densities (Fig. 3D-F).

**Figure 3.**
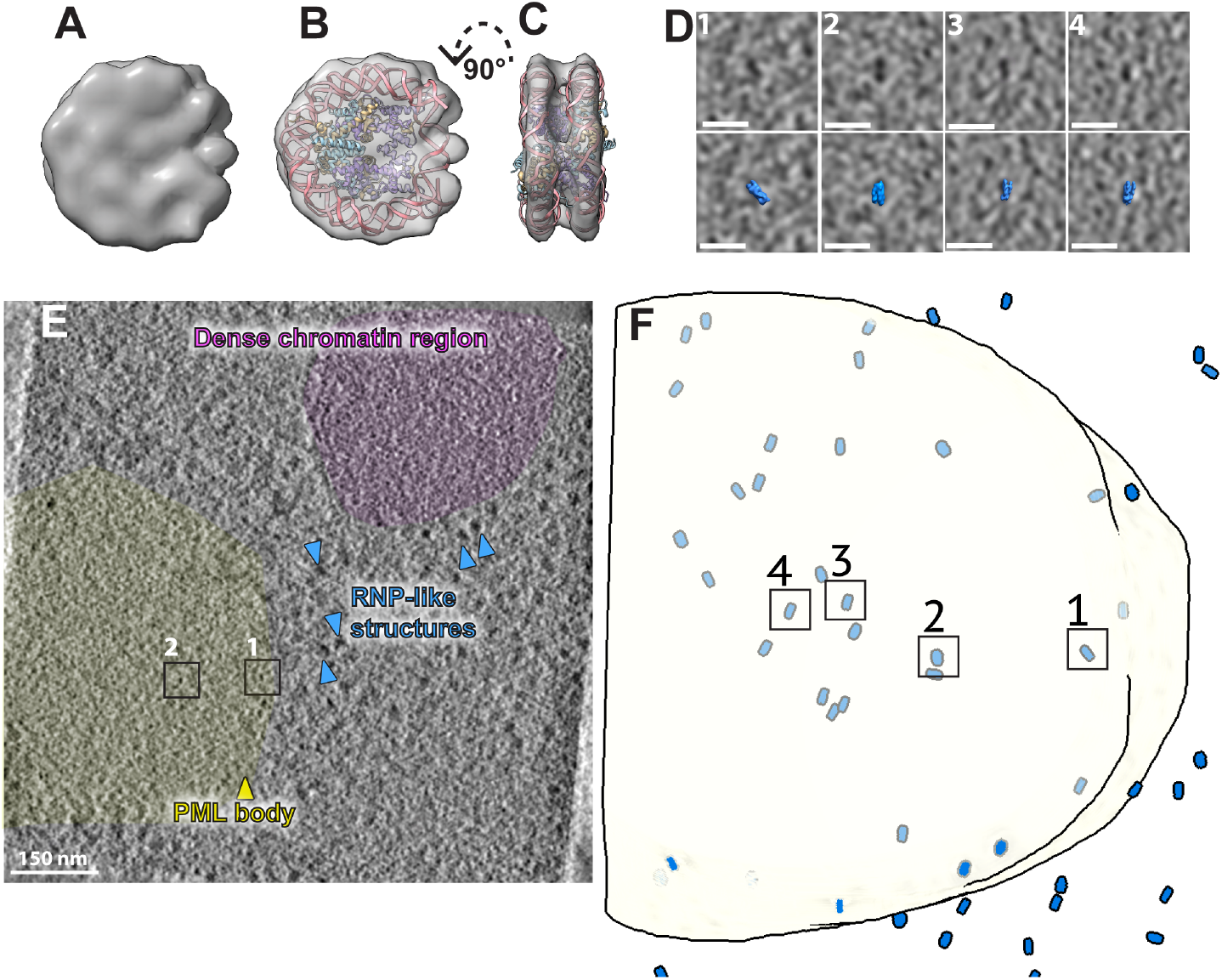
STA of nucleosomes and spatial distribution within PML-I cryo-tomograms. **(A)** STA map of the nucleosome core particle at 19 Å resolution. **(B)** Same as **(A)** rendered transparent with PDB 2CV5 fitted into the density. **(C)** Same as **(B)** rotated 90°; DNA strands are clearly visible. **(D)** Cryo-tomogram slices (top) and corresponding nucleosome particle positions overlaid (bottom, blue) for four representative particles (1–4) indicated in **(F).** Scale bars = 20 nm. **(E)** Cryo-tomogram slice showing the PML-I body environment in TS_15. Dense chromatin region (purple) and RNP-like structures (blue arrowheads) are annotated. Yellow arrowhead indicates the PML-I body; black boxes indicate positions 1 and 2 shown in **(D)**. Scale bar = 150 nm. **(F)** Three-dimensional visual proteomics map showing nucleosome core particle positions mapped inside the segmented PML body mesh (white). Numbered boxes correspond to representative particles shown in **(D)**.

The majority of nucleosomes with or without accessory proteins in cryo-tomograms with segmented PML bodies were located outside (314/390; 80.5%), with 76 particles detected inside (19.5%; Sup. Fig. 15). This fraction is broadly comparable to that of TRiC (14.9% inside), though both contrast with the stronger exclusion of PA28–20S complexes (7.8% inside). The fraction of nucleosomes detected inside varied across tomograms (TS_01: 7.4%; TS_03: 22.6%; TS_08: 23.1%; TS_15: 18.6%; TS_16: 19.0%), reflecting heterogeneity in chromatin content across individual PML-I bodies.

### Columnar trinucleosomes are observed within PML-I bodies

While nucleosomes have been extensively characterised by cryoET, the higher-order chromatin folding long modelled in vitro as two-start zigzag (42) or one-start solenoid (43) fibres has only recently been examined directly in situ, where native chromatin appears as flexible, variable multinucleosome arrangements rather than ordered fibres (44–46). Direct in situ evidence for an ordered columnar trinucleosome arrangement (1) has remained lacking. We therefore sought to determine whether more complex nucleosome structures could be detected in situ within our cryotomograms, using a set of nucleosome-related templates including the open-state columnar trinucleosome structure (PDB: 7V9S; see Methods). On visual inspection of the PyTOM score maps overlaid onto the cryo-tomograms, we noticed that peaks with strong cross-correlation values matched high-contrast densities consistent with multinucleosome assemblies displaying unusual relative orientations (Supplementary Movie 2).

We applied STA with refinement in RELION 4.0 and derived an initial map of a columnar trinucleosome with unusual relative orientation (Fig. 4B, C). We obtained this map by first refining approximately 2,800 particles. The Fourier shell correlation (FSC) curve indicated a resolution of 25 Å at the 0.143 cutoff (Sup. Fig. 17I), with phases randomised beyond 43 Å, indicating that the resolution estimate was not inflated by overfitting to noise. Due to the likely flexibility of the trinucleosome structure, we performed refinement followed by 3D classification focused on the top open nucleosome (Fig. 4D, E; Sup. Fig. 17D-G). This revealed a map at 19 Å resolution with clearly visible major grooves and DNA strands that were not present in our 40 Å low-pass filtered template (Fig. 4D, E; Sup. Fig. 17G, J). We then performed global refinement on the entire trinucleosome to recover the complete structure. In this way, particles were mapped back onto our cryotomograms, revealing high-contrast densities consistent with unusual trinucleosome assemblies (Fig. 4A; Supplementary Movie 1).

**Figure 4.**
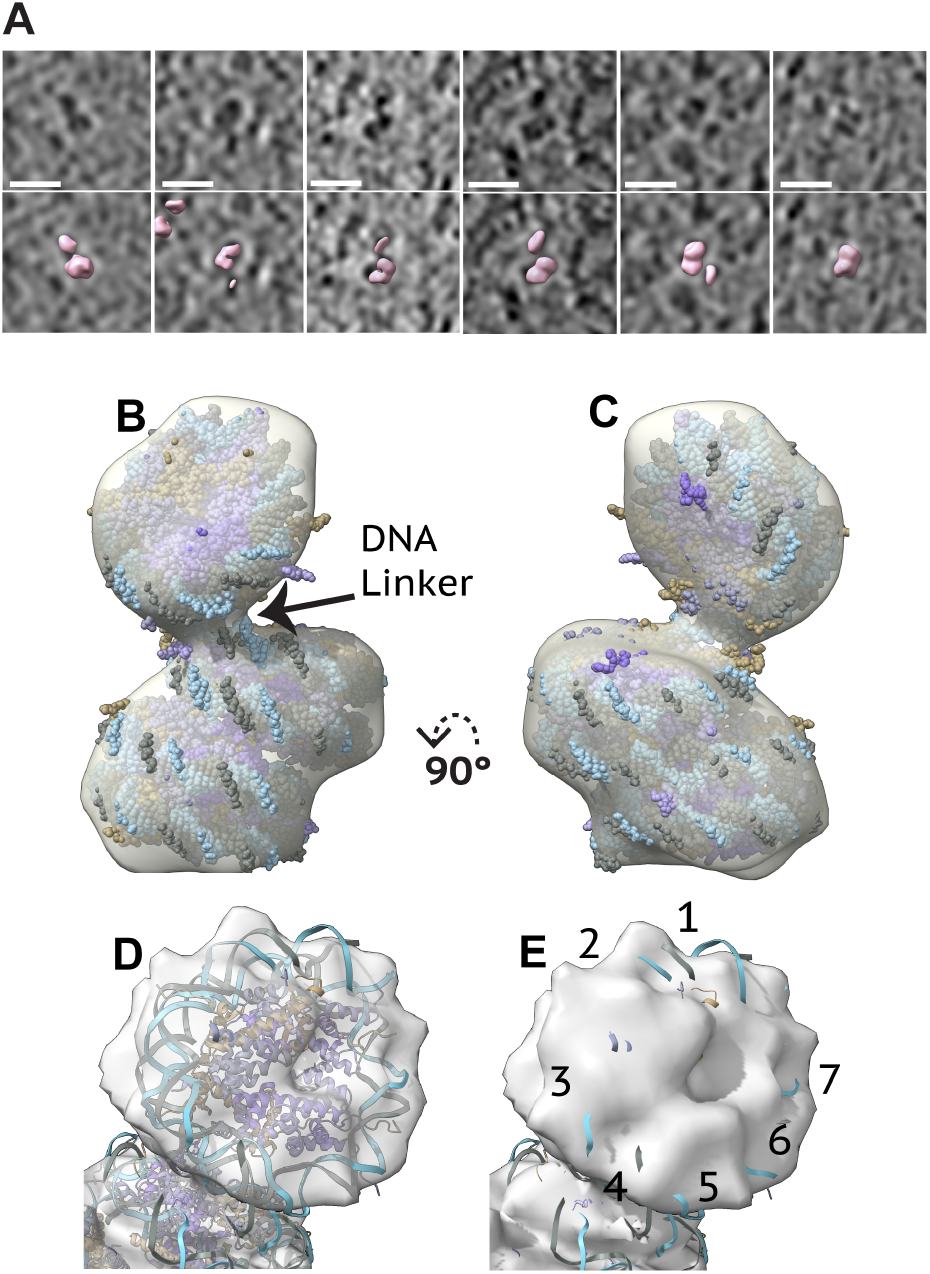
Open trinucleosomes are present within PML-I cryo-tomograms. **(A)** Cryo-tomogram slices (top) and corresponding open trinucleosome particle positions overlaid (bottom, pink) for six representative particles. Scale bar = 20 nm. **(B)** STA map of the open trinucleosome rendered transparent with PDB 7V9S fitted and displayed as a molecular surface. **(C)** Same as **(B)** rotated 90°. **(D)** STA map of the top nucleosome following focused refinement and 3D classification, rendered transparent with PDB 7V9S fitted. **(E)** Same map as in **(D)** of the top nucleosome showing resolved DNA major grooves (numbered 1–7).

To further validate these trinucleosome detections, we next employed a dual-template strategy using both a classical structure and a modified template with an artificial hole introduced into one nucleosome (Sup. Fig. 18A). Independent template matching across all 16 tomograms yielded 2,821 and 1,769 particles from the original and hole templates, respectively. Spatial overlap analysis identified 1,458 particle pairs within 10 nm (51.7% of original-template particles), of which 1,371 (94.0%) showed consistent Y-axis orientations with a mean angular deviation of 8.9°. The consistent orientations between two independently matched particle sets indicate the detections are reproducible rather than template-specific. Critically, STA of the 1,371 validated particles recovered density at the position artificially removed from the hole template (Sup. Fig. 18C, D). Because that density is absent from the input, its recovery is independent evidence that the third nucleosome is genuinely present and not imposed by the reference. Moreover, continuous linker DNA density connecting adjacent nucleosomes is resolved in the trinucleosome map (Fig. 4B), excluding the alternative that these detections represent three independently positioned mononucleosomes in incidental proximity. Together with the major-groove and DNA features that emerged at 19 Å but were absent from the 40 Å template (Fig. 4D, E; Sup. Fig. 17J), these results confirm the in situ presence of columnar open trinucleosome assemblies within intact cell nuclei, consistent with the architecture defined by in vitro cryo-EM (1) but observed here directly for the first time.

Mapping both particle sets onto the five segmented PML-I body meshes showed that nucleosomes (76/390; 19.5%) and trinucleosomes (143/763; 18.7%) were detected inside PML-I body interiors at comparable rates (Sup. Fig. 15, 16). We then sought to determine even higher-order nucleosome arrangements. To investigate the spatial co-organisation of nucleosome and trinucleosome assemblies across the full dataset, we performed joint DBSCAN clustering of all 4,045 detected particles (1,758 nucleosomes across 15 tomograms; 2,287 trinucleosomes across 13 tomograms). This combined analysis revealed that 3,756 particles (92.9%) were organised into 136 discrete spatial clusters, with cluster sizes ranging from 3 to 524 particles (Sup. Fig. 19). Notably, 111 of the 136 clusters (81.6%) were classified as mixed, containing both nucleosome and trinucleosome particles, demonstrating that mono- and trinucleosome assemblies are co-localised within the same spatial domains rather than segregated into distinct zones. In the five cryo-tomograms with available PML-I body segmentations (TS_01, TS_03, TS_08, TS_15, TS_16), the dominant clusters were spatially coincident with the PML body interior or its immediate periphery (Fig. 5A-C; Sup. Fig. 19A–E). The largest clusters in TS_08 and TS_15 comprised 144 and 169 particles, respectively, and co-localised with the PML-I body boundary, consistent with the dense chromatin observed in these regions (Sup. Fig. 19F-G). Together, these data indicated that nucleosome and trinucleosome particles are organised into discrete, spatially coherent clusters, with the largest assemblies preferentially associated with PML bodies. To test whether this spatial organisation exceeds random expectation, we applied second-order Ripley’s L function analysis (47) across all cryo-tomograms, comparing observed inter-particle distances against 200 Monte Carlo simulations of complete spatial randomness (CSR) within the same volume. Statistically significant clustering was detected in all 15 tomograms containing nucleosome detections, in all 13 tomograms containing trinucleosome detections, and significant spatial co-clustering between the two particle types was detected in all 13 tomograms where both were present (p < 0.05; Sup. Fig. 20). The clustering was most pronounced at spatial scales of 34–77 nm, consistent with the physical extent of short chromatin fibre segments containing multinucleosome or trinucleosome assemblies.

**Figure 5.**
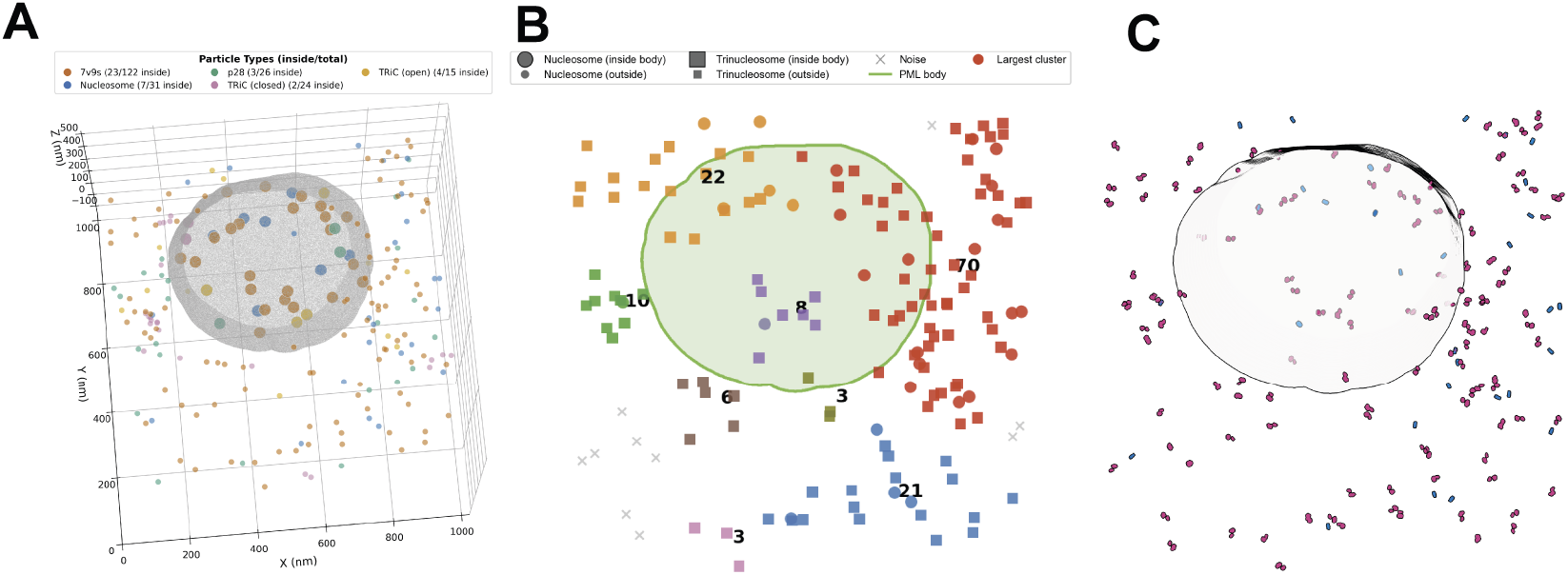
Spatial distribution of macromolecular complexes within PML-I bodies. **(A)** Three-dimensional distribution of all detected particle types relative to the PML-I body mesh in tomogram TS_03. Particle types are colour-coded as indicated. Large markers denote particles localised within the PML body interior (as determined by mesh containment analysis); small markers denote particles outside. The PML body surface is rendered as a Fresnel-shaded mesh. **(B)** Combined DBSCAN spatial clustering of nucleosome (circles) and trinucleosome (squares) particles in TS_03 (eps = 100 bin4 pixels (∼99 nm), min_samples = 3). Clusters are colour-coded; the largest cluster is shown in red, with cluster sizes indicated at centroids. Large markers denote particles inside the PML body; small markers denote particles outside. The PML body boundary is shown as a green contour. Noise particles are shown as crosses. **(C)** Trinucleosome particles mapped back onto the PML body shell in TS_03, displayed as STAs at their refined positions and orientations. Trinucleosomes are shown in magenta; nucleosomes in blue. The PML body shell is shown in white.

This detection of nucleosome and columnar trinucleosome assemblies provides the first direct molecular-resolution visualisation of nucleosomal chromatin within PML-I body interiors. The predominance of mixed mono/trinucleosome clusters (81.6% of 136 clusters) indicates that chromatin within and around PML bodies does not segregate by nucleosome conformation but instead forms compositionally heterogeneous assemblies, consistent with a dynamic, interconverting chromatin network. The selective admission of these chromatin assemblies (19.5% inside), alongside a broadly comparable interior fraction of TRiC (14.9%) but stronger exclusion of PA28-capped proteasomes (7.8%), indicates that the PML-I body interior partitions macromolecular complexes by size.

## Discussion

Our cryo-ET and cryo-SR-CLEM measurements now provide a direct view of the molecular interior of intact PML-I bodies in situ. Columnar trinucleosome assemblies and nucleosomes are detected here, providing the first direct molecular-resolution visualisation of nucleosomal chromatin within PML-I body interiors. Together, these observations reveal that the PML-I body interior is a fibrous network whose pores may selectively partition macromolecular complexes by size. This architecture is consistent with predictions of the phase-separation-coupled-to-percolation (PSCP) framework for multivalent biomolecular condensates (48–51), reframing the PML body from a featureless protein condensate into a chromatin-permissive, structurally heterogeneous compartment. Our data now suggest that one function of this framework is to create a ‘chromatographic’, porous core that can drive physical separation of contents for function-associated selection.

The stronger exclusion of PA28-capped 20S proteasomes (7.8% inside) supports size-selective partitioning by the porous mesh, while TRiC is present at an interior fraction comparable to chromatin (14.9% versus 19.5%), with post-translational modifications likely contributing further to compositional specificity.

The combined power of cryo-ET and cryo-SR-CLEM resolves nuclear condensate interiors, enabling the molecular dissection of PML-I body composition. The same workflow should be tractable for other shell-core or multiphase nuclear condensates such as paraspeckles and nucleoli. More broadly, our integrated cryo-ET and cryo-SR-CLEM pipeline provides a generalisable framework for investigating condensate architecture at molecular resolution.

As well as informing the physical organisation of PML bodies, our TRiC data provide mechanistic insight into nuclear protein folding. We resolve both open and closed TRiC conformations in the nucleus, consistent with an ongoing chaperonin duty cycle within the nuclear environment (39). The presence of TRiC inside at a minority fraction (14.9% of all TRiC), together with conformation-dependent clustering of closed-state TRiC in the adjacent nucleoplasm, indicates that TRiC-mediated protein folding occurs mainly around, rather than within, PML-I bodies. This spatial segregation suggests that PML bodies and the protein folding machinery occupy distinct but neighbouring nuclear compartments.

Vault complexes are classically regarded as cytoplasmic, and to our knowledge have not previously been visualised at structural resolution within the nucleus. Our cryo-ET detection of these in the nucleus was corroborated by immunofluorescence against the major vault protein. Our observation of these at the periphery of PML bodies may reflect a role in nucleocytoplasmic trafficking of associated factors, given previously established roles for vaults in transport to the nucleus (52), drug resistance (53) and in innate immune signalling (54).

Our detection of membrane vesicles within and adjacent to PML bodies is similarly unexpected, given that PML bodies are classically defined as membrane-less compartments. We propose that these membrane structures most likely represent remnants of nuclear envelope reformation following mitosis, in which membrane fragments become transiently enclosed within the reforming nucleus before being resolved. Characterising these membrane-associated structures in additional cell types and conditions will be an important avenue for future work.

A central finding of this study is the surprising detection of nucleosomes and trinucleosome assemblies within PML-I body interiors. Notably, previous electron spectroscopic imaging (ESI) data suggested that PML body cores lack detectable nucleic acids in chemically fixed samples (28). The absence of signal in that study likely reflects the local organisation of internalised chromatin, which we now observe, may fall below the sensitivity threshold of ESI, potentially also combined with displacement during chemical fixation (55).

Our precise in situ structural observations are consistent with a growing body of evidence that suggests that specific PML body subtypes incorporate DNA (19, 20, 22–24), and now extends this catalogue to importantly include nucleosomal chromatin and trinucleosomes. Interestingly, the most successful trinucleosome template used here was derived from a structure of human telomeric chromatin (1), that also bears a columnar arrangement of nucleosomes in an open conformation. This is characterised by a short nucleosome repeat length. The assemblies we have detected within PML-I bodies also display this open conformation and we also observed an accessory density. Further work will be required to resolve these assemblies at higher resolution and to determine which accessory proteins (e.g. TRF-1) can be resolved within the detected structures.

This and the prevalence of this open-state conformation within and near PML bodies suggest that the PML body chromatin is accessible to chromatin-binding factors. The open state in telomeric chromatin has been suggested to expose the H2A-H2B acidic patch for interaction with chromatin remodellers, SIRT6 and DNA damage response proteins, without requiring decompaction of the surrounding heterochromatin (1). PML bodies have established roles in the DNA damage response and in chromatin remodelling at specific loci, including telomere maintenance in ALT-positive cells (17, 22, 23), and the open trinucleosome assemblies observed here are consistent with this regulatory context. Furthermore, PML bodies have been reported to associate with pericentromeric satellite DNA during G2 phase (24), raising the possibility that a subset of the columnar trinucleosome assemblies detected here represents pericentromeric rather than telomeric chromatin. Notably, like the telomeric repeat sequence used to determine the columnar open trinucleosome structure (1), pericentromeric satellite DNA is characterised by highly repetitive sequences, suggesting that this open conformation may be a general structural feature of repetitive chromatin. As our HFF cultures were not cell-cycle synchronised, the precise chromatin context of these assemblies remains to be determined.

Finally, our network connectivity analysis reveals that the interior mesh is consistently denser and more highly connected than the surrounding nucleoplasm (Pearson r = 0.52, 0.56 and 0.31 for TS_01, TS_03 and TS_08, Sup. Fig. 2D). PML bodies are well established to concentrate hundreds of small client proteins through SUMO-SIM interactions (5, 6), and soluble protein regulators freely partition into the body interior. Our observation now extends this picture by placing a structural constraint at the other extreme of the size range, where PA28 single-capped proteasomes are most strongly excluded (7.8% of particles localised within the body) and TRiC chaperonin complexes are present at a minority fraction comparable to chromatin (14.9%), while nucleosomes and trinucleosomes access the interior (19.5%). Our study now provides structural evidence for a selective permeability not previously observed at molecular resolution in PML bodies, and suggests a physical mechanism in which the mesh geometry may accommodate small client proteins and flexible chromatin fibres while restricting large globular complexes of ∼17–20 nm diameter. Only ∼28% of pores exceed the ∼17 nm diameter of a TRiC barrel and ∼15% the ∼20 nm diameter of a PA28-capped proteasome, whereas ∼79% readily accommodate the ∼11 nm diameter of a nucleosome, providing a geometric rationale consistent with the observed preferential accommodation of chromatin over large globular complexes. Porous meshwork architectures have recently been resolved by cryo-ET in the pericentriolar material of C. elegans centrosomes (36) and in the polar organiser PopZ condensate in bacteria (56, 57), and selective permeability has been proposed but not yet observed for stress granules (58). Together, these examples, alongside our determination of a network structure in PML bodies, suggest that a percolated, porous meshwork may be a general architectural feature of biomolecular condensates, which may operate in a manner reminiscent of size-exclusion chromatography (SEC). Notably, PML (TRIM19) and TRIM5α are closely related TRIM family members, and TRIM5α similarly forms organised cytoplasmic bodies directly visualised by cryo-ET (33); this raises the further possibility that such internal network organisation (and ‘intracellular SEC’) is a general mechanism and a conserved property of TRIM family condensates.

## Limitations

Several constraints of our approach should be acknowledged. First, our cryo-lamellae are 134 to 230 nm thick sections through PML-I bodies, which can reach ∼800 nm in diameter, so each cryo-tomogram captures only a thin slab of an individual body rather than its full three-dimensional volume. Our visual-proteomics statistics (n = 5 segmented tomograms) therefore describe the particle content of these slabs, and absolute particle counts inside the body should be interpreted as per-slab rather than per-body totals. Second, template matching is intrinsically biased toward the templates searched, and computational 3D classification during sub-tomogram averaging can lose a substantial fraction of true-positive particles (59). Our final particle counts (389 PA28, 297 closed and 276 open TRiC, and 1,371 validated trinucleosomes) should therefore be interpreted as lower bounds on the abundance of each complex. Third, resolving finer features, for example accessory proteins bound to the trinucleosome, will require larger particle sets and multi-particle tilt-series refinement strategies. Finally, our trinucleosome template (PDB 7V9S) was derived from a reconstituted human telomeric chromatin array, so our in situ detections are consistent with this open-state columnar arrangement but do not by themselves establish telomeric identity or exclude similar assemblies at non-telomeric loci.

## Acknowledgements

We thank the Beatson Advanced Imaging Resource (RRID: SCR_023875) for imaging support. Cryo-CLEM and cryo-ET data collection were performed at the Multi-User CryoEM Facility at CSSB, supported by the University of Hamburg and DFG grant numbers INST 152/772-1, 152/774-1 and 152/776-1 FUGG; and at the Rosalind Franklin Institute; we thank Thomas Glen and Matthew Case for support with cryo-FIB milling and cryo-ET data collection respectively, facilitated through the Electrifying Life Sciences programme (Wellcome Trust 220526/Z/20/Z). We thank Maud Dumoux for helpful discussions.

This work was supported by core grants to the CRUK Scotland Institute (A17196 and A31287; ROR 03pv69j64). S.D.C. was supported by a UK Research and Innovation Future Leaders Fellowship (MR/W010690/1), a Royal Society Research Grant (RGS\R1\231494) and a Medical Research Council programme grant (MR/Z504464/1). C.B. was supported by Medical Research Council grant MC_UU_00034/2. R.K. was supported by a Volkswagen Foundation Freigeist Fellowship (91671; 91671-2) and by the Deutsche Forschungsgemeinschaft (DFG) grant number 496128632. The Rosalind Franklin Institute is funded by the UK Research and Innovation through the Engineering and Physical Sciences Research Council (UKRI-EPSRC); Next Generation Chemistry at the Rosalind Franklin Institute has been supported by grants EP/V011 359/1, EP/T012 021/1, EP/X527 245/1. S.D.C. held a sabbatical at the Rosalind Franklin Institute supported by the Electrifying Life Sciences programme (Wellcome Trust 220526/Z/20/Z).

## Author contributions

S.D.C. conceived and supervised the project. V.P., R.K. and S.D.C. collected the cryo-ET data. I.H. and P.T. performed 3D-SIM. R.K., J.F. and V.P. performed cryo-SR-CLEM. All authors contributed to data analysis, manuscript writing and editing.

## Data, code, and materials availability

Sub-tomogram averaging maps have been deposited in the Electron Microscopy Data Bank (EMDB) under accession codes EMD57924 (TRiC closed, 22 Å), EMD-58330 (PA28 single-capped, 29 Å), EMD-57913 (nucleosome, 19 Å), EMD-57971 (trinucleosome focused refinement, 19 Å), and EMD-57970 (trinucleosome full map, 25 Å). Raw cryo-tomographic data have been deposited in the Electron Microscopy Public Image Archive (EMPIAR) under accession code [EMPIAR-XXXXX]. Analysis scripts used in this study are available in four GitHub repositories: PML body internal network analysis (https://github.com/StephenC07/pml-body-network-analysis); a napari-based PyTOM CC-score threshold viewer (https://github.com/StephenC07/pytom-cc-threshold-viewer); mesh-based PML body visual proteomics (https://github.com/StephenC07/pml-body-visual-proteomics); and the TRiC cluster pair-orientation classification (https://github.com/StephenC07/tric-cluster-orientation-analysis). The doxycycline-inducible PML-I human foreskin fibroblast cell line is available upon reasonable request from the corresponding author. Supplementary Movie 1 is available on Zenodo at https://zenodo.org/records/20476347.

## Supplementary figure legends

*Main figure legends (Figures 1–5) accompany their respective figures*.

**Supplementary Figure 1.**
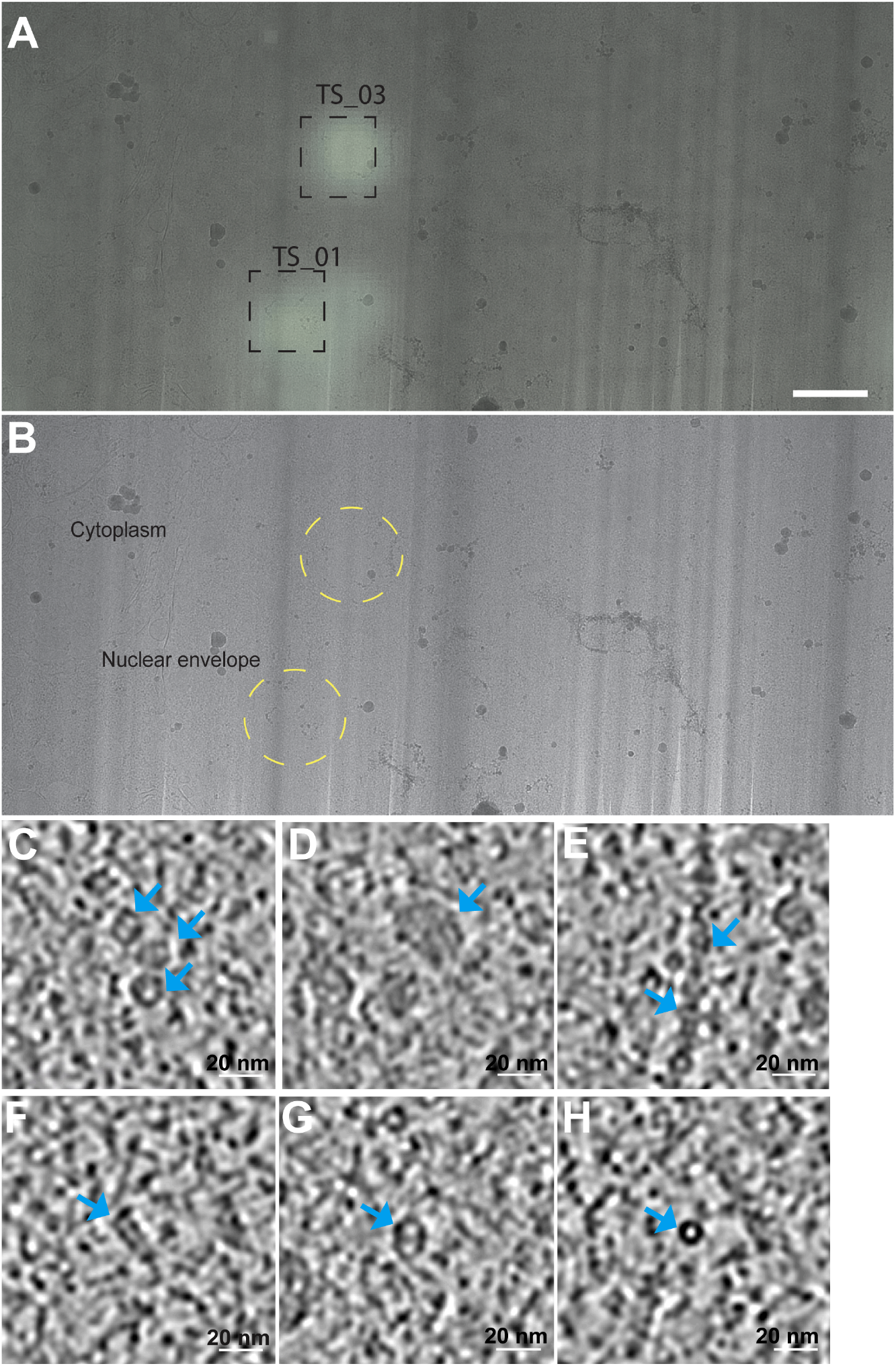
Cryo-CLEM overview and representative high-contrast densities in eYFP-PML-I body cryo-tomograms. **(A)** Cryo-fluorescence image of a vitrified lamella showing two eYFP-PML-I bodies (TS_01, TS_03; dashed boxes). Scale bar = 1 µm. **(B)** TEM overview of the same lamella with PML body positions indicated by yellow dashed circles; cytoplasm and nuclear envelope are labelled. **(C–H)** Representative cryo-tomogram slices illustrating the morphological diversity of high-contrast densities observed within and adjacent to PML body interiors: capsules **(C)**, granules **(D)**, filaments **(E)**, cylindrical structures **(F)**, large barrels **(G)**, and rings **(H)**. Scale bars = 20 nm.

**Supplementary Figure 2.**
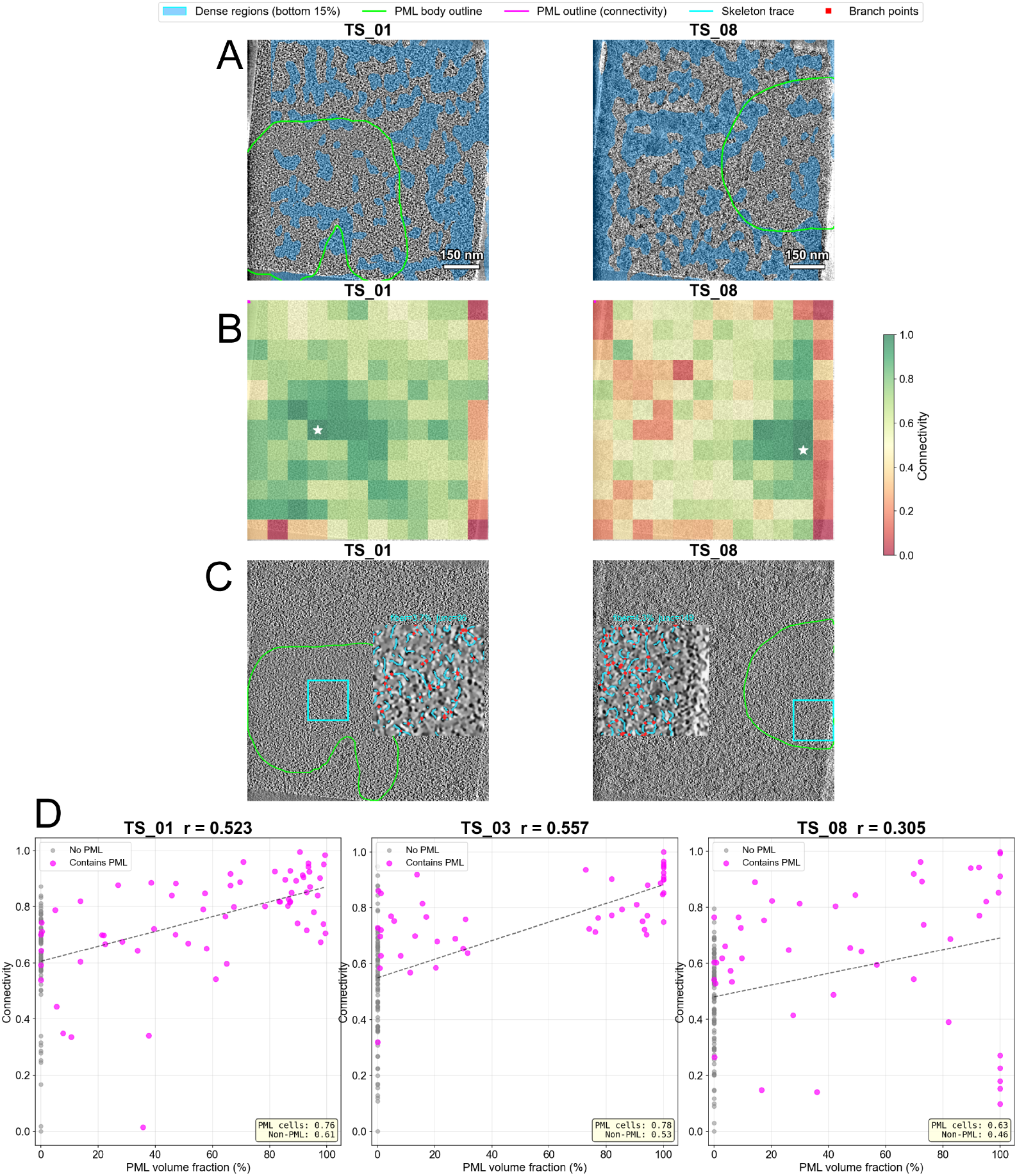
eYFP-PML-I body network analysis. **(A)** Z-projections of cryo-tomogram slices (TS_01, TS_08) showing high-contrast dense regions (cyan, bottom 15% intensity) relative to the segmented PML body (green outline). Dense regions are spatially excluded from the PML body interior in both tomograms. **(B)** Skeleton connectivity heatmaps (12×12 grid) overlaid on mean-intensity projections of TS_01 and TS_08. Connectivity score (0–1, red–green) was computed from black top-hat segmentation followed by 2D skeletonization of each slice, combining fibre density (60%) and branch point density (40%). White star indicates the most connected grid cell. Higher connectivity is observed within and adjacent to the PML body in both cryo-tomograms. **(C)** Representative cryo-tomogram slices (TS_01, TS_08) with skeleton trace insets. Cyan box indicates the most connected grid region; inset shows the denoised crop with skeleton overlay (cyan) and branch points (red). **(D)** Quantification of skeleton connectivity as a function of eYFP-PML-I body volume fraction per grid cell. Each point represents one grid cell; magenta points contain PML body, grey points do not. Dashed line shows linear fit. Positive correlations (r = 0.52, 0.56, 0.31) confirm that network density increases with PML body occupancy.

**Supplementary Figure 3.**
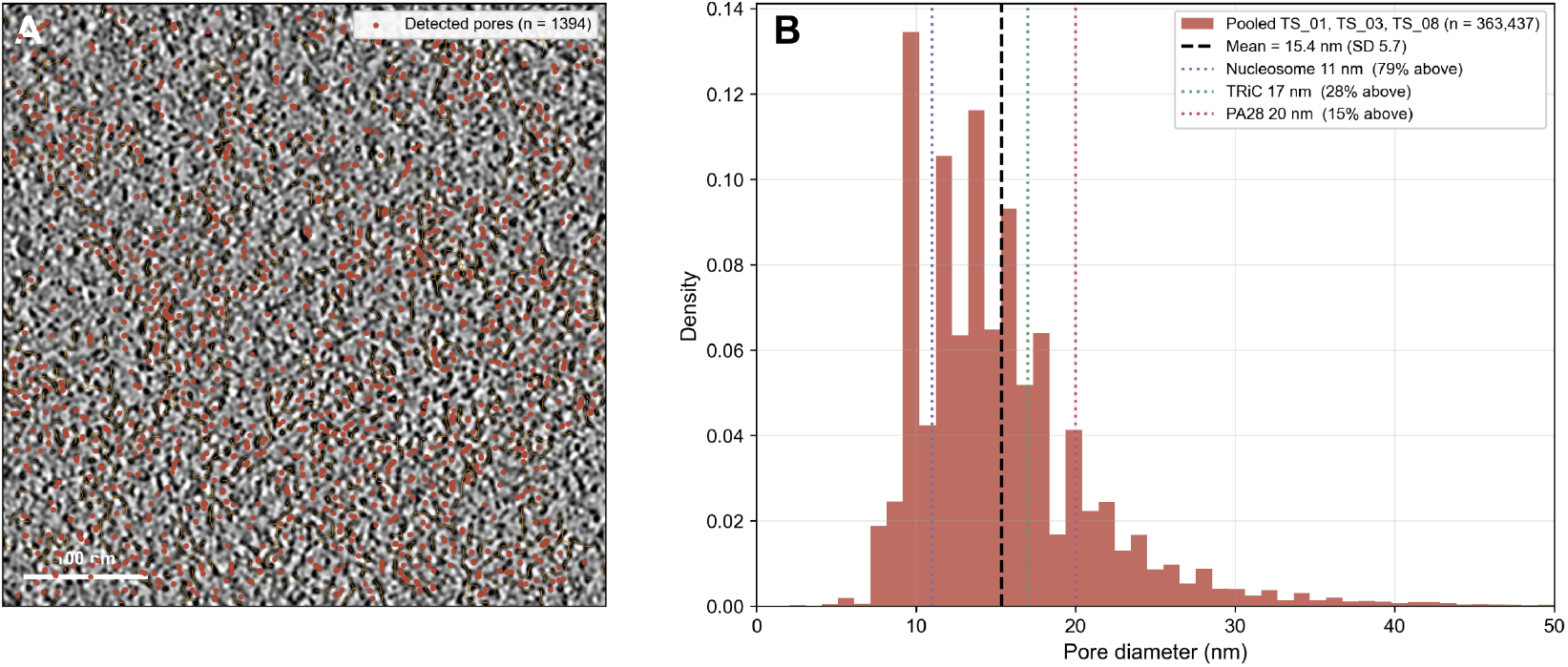
Pore-size analysis of the eYFP-PML-I body interior. **(A)** Representative cryo-tomogram slice (TS_03, 512 by 512 pixel crop) with the fibre skeleton overlaid in orange and detected pore centres in red. Scale bar = 100 nm. **(B)** Pooled pore-diameter distribution from TS_01, TS_03 and TS_08 (n = 363,437). Mean = 15.4 nm, standard deviation 5.7 nm (black dashed line). Dotted lines mark nucleosome (11 nm), TRiC (17 nm) and PA28 (20 nm) diameters. 79 percent of pores accommodate a nucleosome, 28 percent a TRiC, 15 percent a PA28.

**Supplementary Figure 4.**
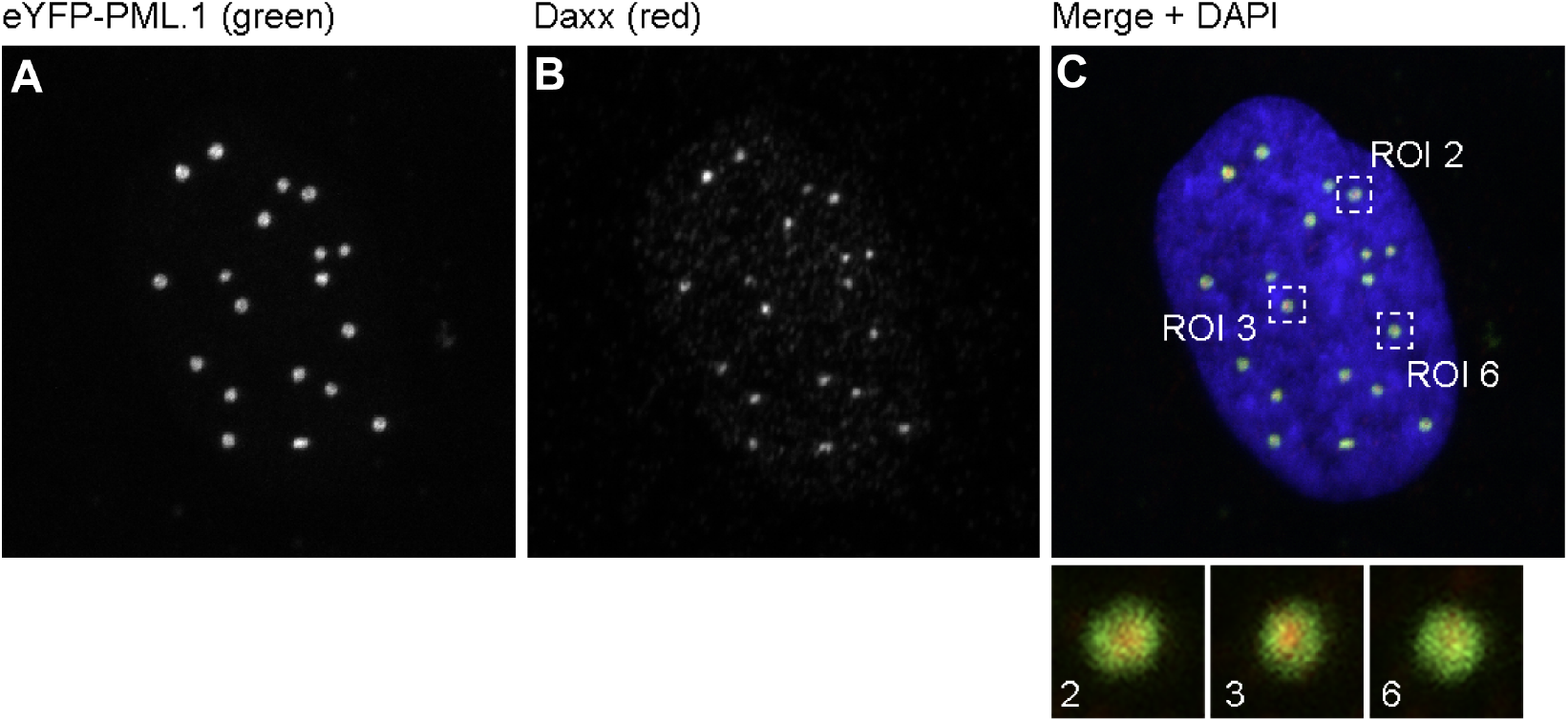
Daxx immunofluorescence as a positive control for PML body colocalisation. **(A)** eYFP-PML-I channel (green) showing eYFP-PML-I nuclear bodies. **(B)** Daxx immunofluorescence (red), a known PML body component. **(C)** Merged image with DAPI (blue) showing colocalisation of eYFP-PML-I and Daxx signals. Insets show magnified ROIs 2, 3, and 6, confirming Daxx colocalisation with PML bodies.

**Supplementary Figure 5.**
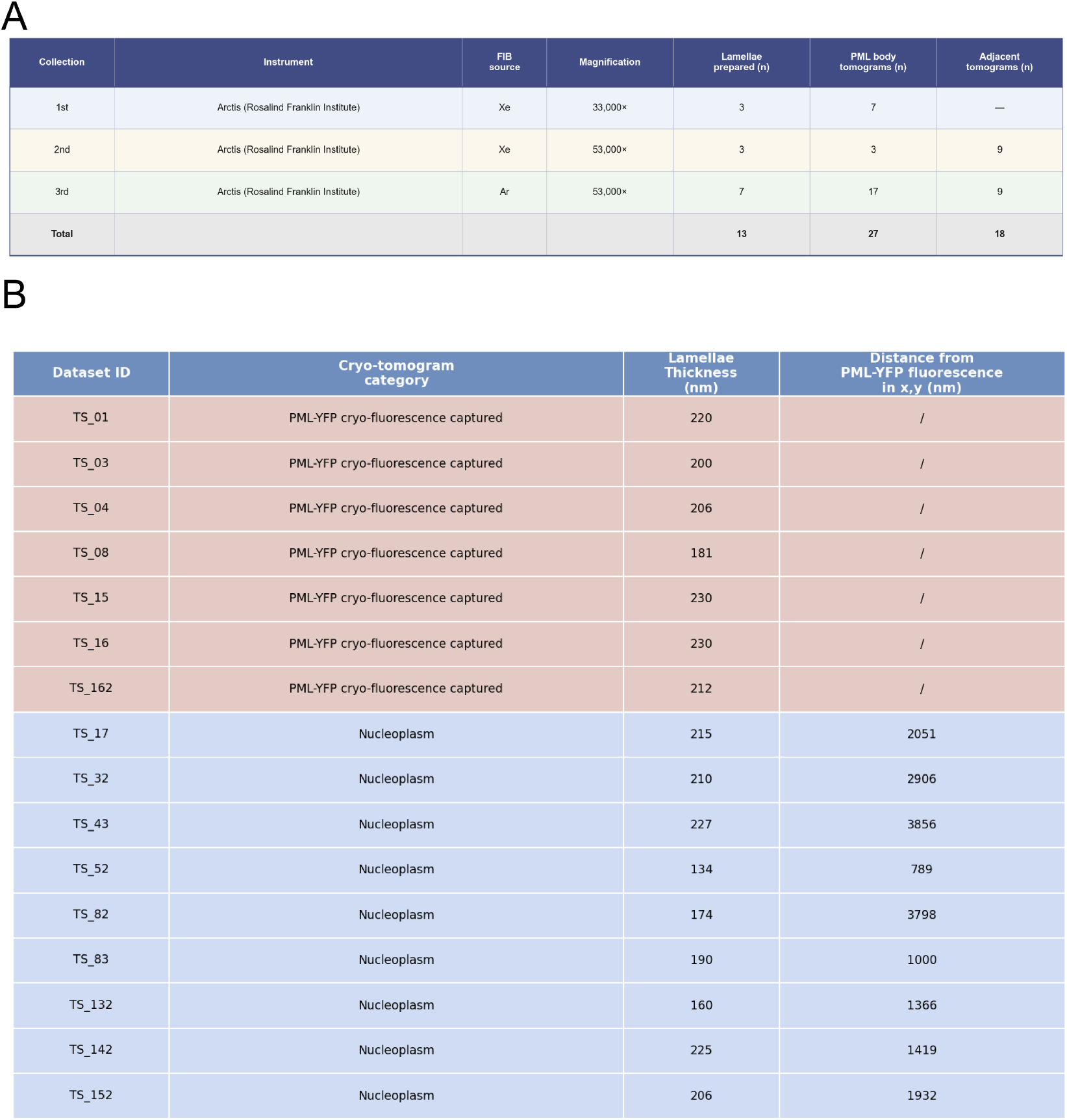
Cryo-pFIB data collection and cryo-tomogram dataset summary. **(A)** Summary of cryo-pFIB lamella data collection across three sessions, detailing the instrument, pFIB plasma source, magnification, number of lamellae prepared, and PML body and adjacent nuclear tomograms collected per session. **(B)** Table summarising the 16 cryo-tomogram datasets selected for analysis, with eYFP-PML-I cryo-fluorescence captured cryo-tomograms listed first (pink). Columns indicate the dataset ID, cryo-tomogram category, lamella thickness (nm), and distance from eYFP-PML-I fluorescence signal in x,y (nm). “/” denotes tomograms in which an eYFP-PML-I body was directly captured within the field of view. Nucleoplasm datasets (blue) were collected from nuclear regions not containing a captured eYFP-PML-I body.

**Supplementary Figure 6.**
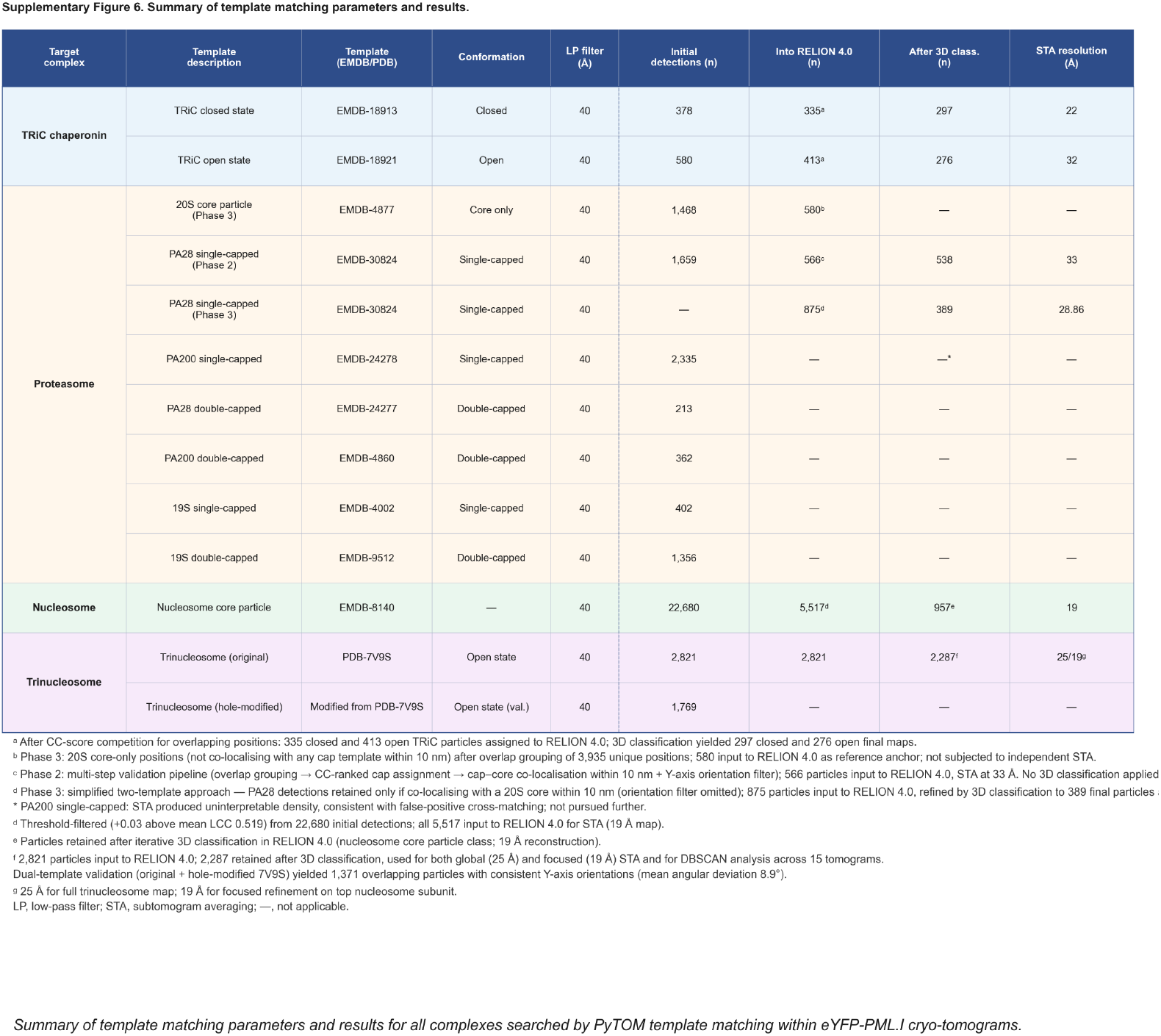
Summary of template matching parameters and results. Summary of all template matching searches performed in PyTOM and subsequent STA workflow in RELION 4.0, including template descriptions, initial detection counts, particles carried into RELION 4.0, particles retained after 3D classification, and final STA resolution.

**Supplementary Figure 7.**
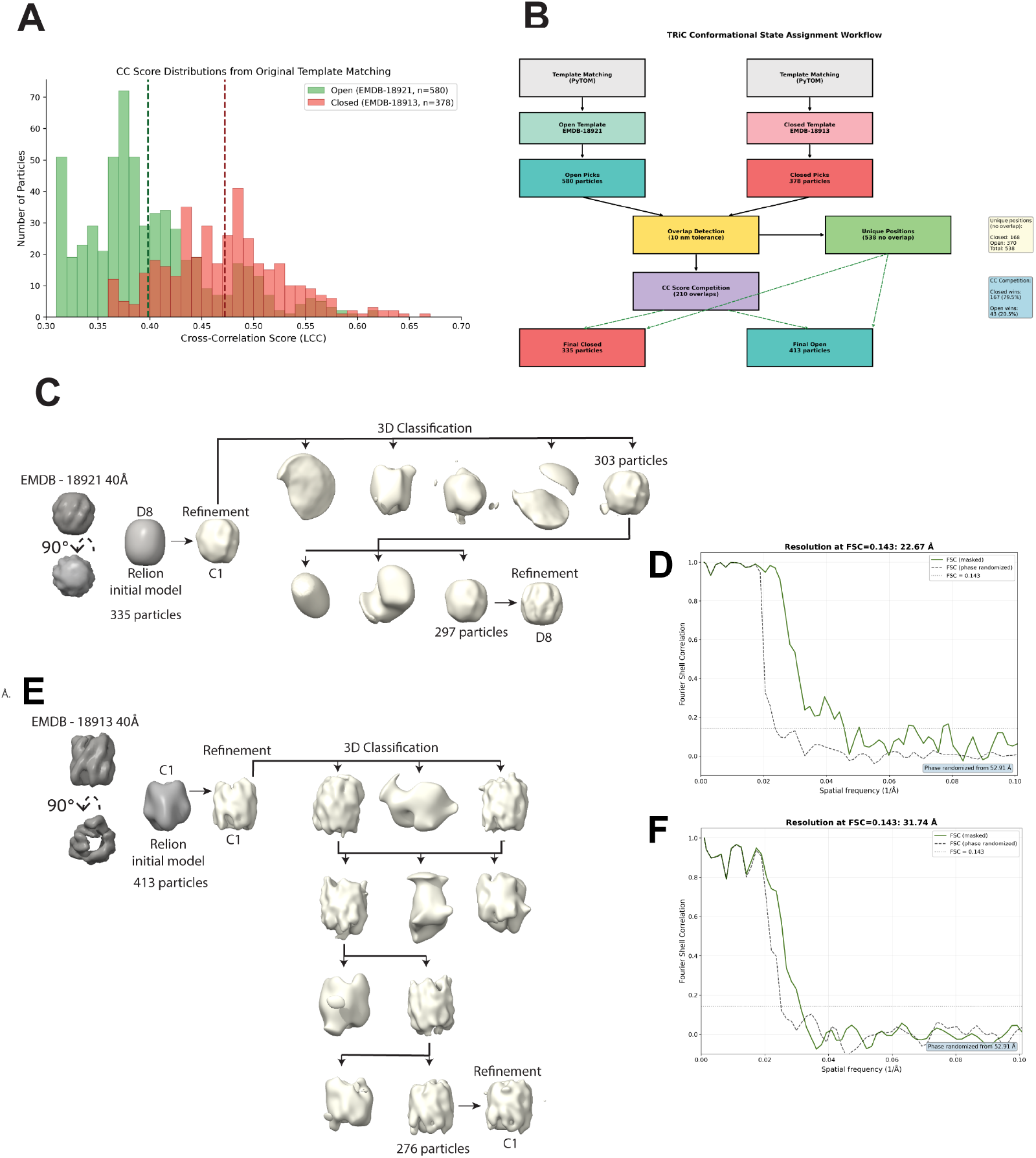
Template matching and STA workflow of open and closed TRiC. **(A)** CC score distributions open (green, EMDB-18921, n = 580) and closed (pink, EMDB-18913, n = 378) TRiC templates, **(B)** Flowchart illustrating the TRiC conformational state assignment pipeline. Template matching yielded 580 open and 378 closed particle candidates. Overlap detection with a 10 nm tolerance identified 538 unique positions. The remaining 210 overlapping positions were resolved by CC score competition, in which closed TRiC won 167 competitions (79.5%), yielding final sets of 335 closed and 413 open TRiC particles. **(C)** STA workflow for closed TRiC (EMDB-18913), starting from 335 particles. C1 refinement followed by two rounds of 3D classification with 5 and 3 classes respectively. D8 refinement of 297 filtered particles yielded the final reconstruction. **(D)** FSC curve for the closed TRiC STA. Resolution = 22 Å at FSC = 0.143. **(E)** STA workflow for open TRiC (EMDB-18921), starting from 413 particles. Iterative 3D classification, followed by refinement in C1 of 276 filtered particles. **(F)** FSC curve for the open TRiC STA. Resolution = 32 Å at FSC = 0.143.

**Supplementary Figure 8.**
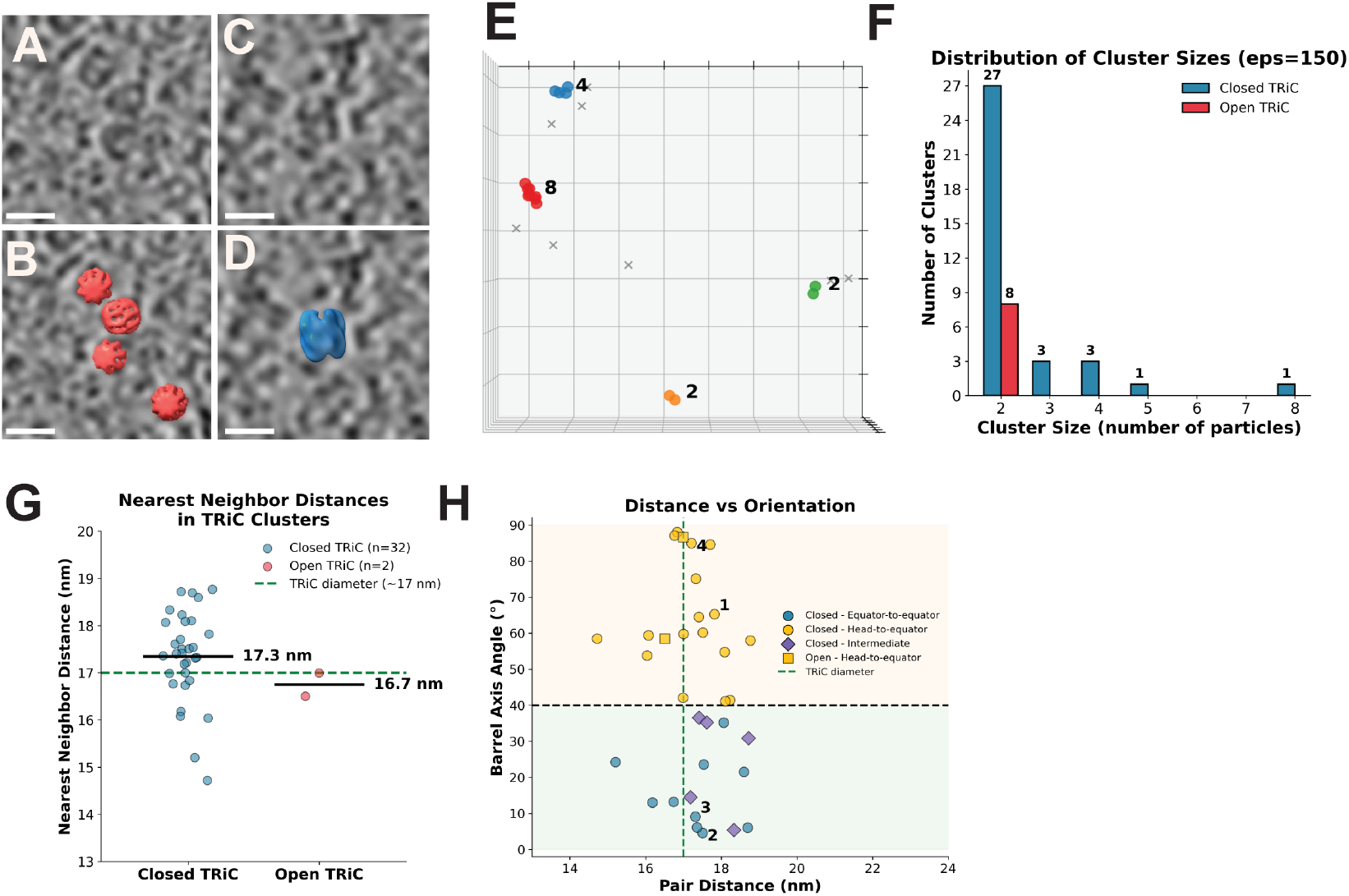
TRiC chaperonin cluster organisation and spatial analysis. **(A)** Cryo-tomogram slice showing closed TRiC cluster densities. **(B)** Closed TRiC STAs mapped back onto the tomogram (red). **(C)** Cryo-tomogram slice showing an open TRiC cluster density. **(D)** Open TRiC STA mapped back onto the cryo-tomogram (blue). Scale bars = 20 nm. **(E)** Two-dimensional DBSCAN cluster map showing spatial positions of closed TRiC particles, with cluster sizes indicated. **(F)** Distribution of TRiC cluster sizes determined by DBSCAN (eps ≈ 37 nm) for closed (blue) and open (red) TRiC. **(G)** Nearest-neighbour distances within TRiC clusters for closed (n=32, mean 17.3 nm) and open (n=2, mean 16.7 nm). Dashed line indicates TRiC diameter (∼17 nm). **(H)** Distance versus orientation scatter plot showing barrel axis angle against pair distance for TRiC contact geometries: closed equator-to-equator (blue circles), closed head-to-equator (yellow circles), closed intermediate (purple diamonds), and open head-to-equator (yellow squares). Dashed lines indicate the TRiC diameter and the 40° barrel-axis threshold. Numbered points (1–4) mark example pairs displayed in Fig. 2V–W.

**Supplementary Figure 9.**
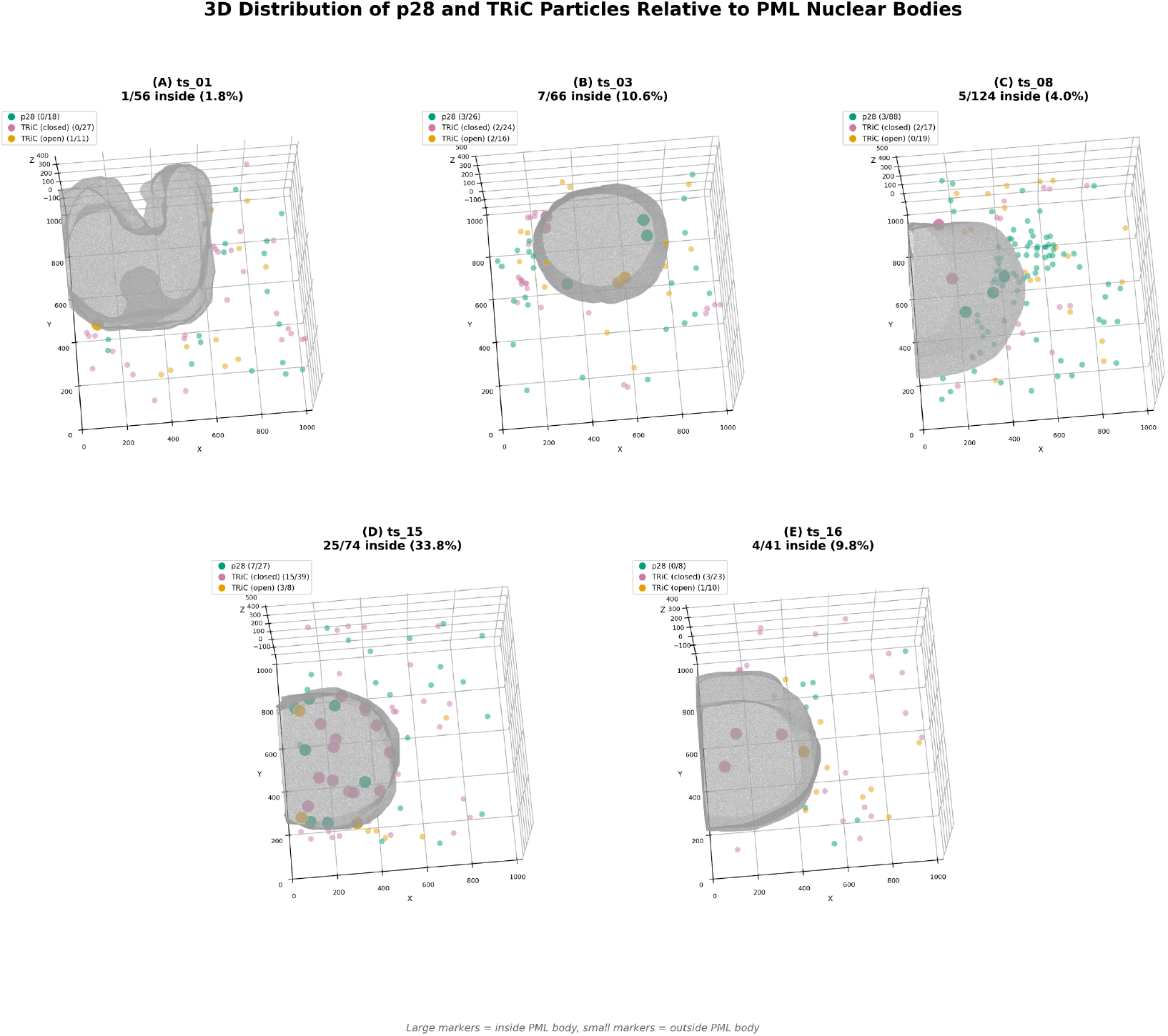
3D distribution of PA28 and TRiC particles relative to PML-I bodies. **(A-E)** 3D renderings of PML body segmentations from five tomograms (A, TS_01; B, TS_03; C, TS_08; D, TS_15; E, TS_16), showing the spatial distribution of PA28 (green), TRiC closed (pink), and TRiC open (orange) particles. PML body surface meshes (grey, semi-transparent) were segmented in Amira and rendered using isosurface extraction via marching cubes. Large spheres indicate particles classified as inside the PML body, small spheres indicate particles outside. Inside/outside classification was performed using ray-casting on the triangulated mesh surface. Coordinates were scaled to bin4 pixel space (divide by 4) to match the segmentation volumes. Z-axis exaggeration (1.2x) and tilt corrections (25 deg X-tilt, 10 deg Y-tilt) were applied to better visualise the plots in 3D.

**Supplementary Figure 10.**
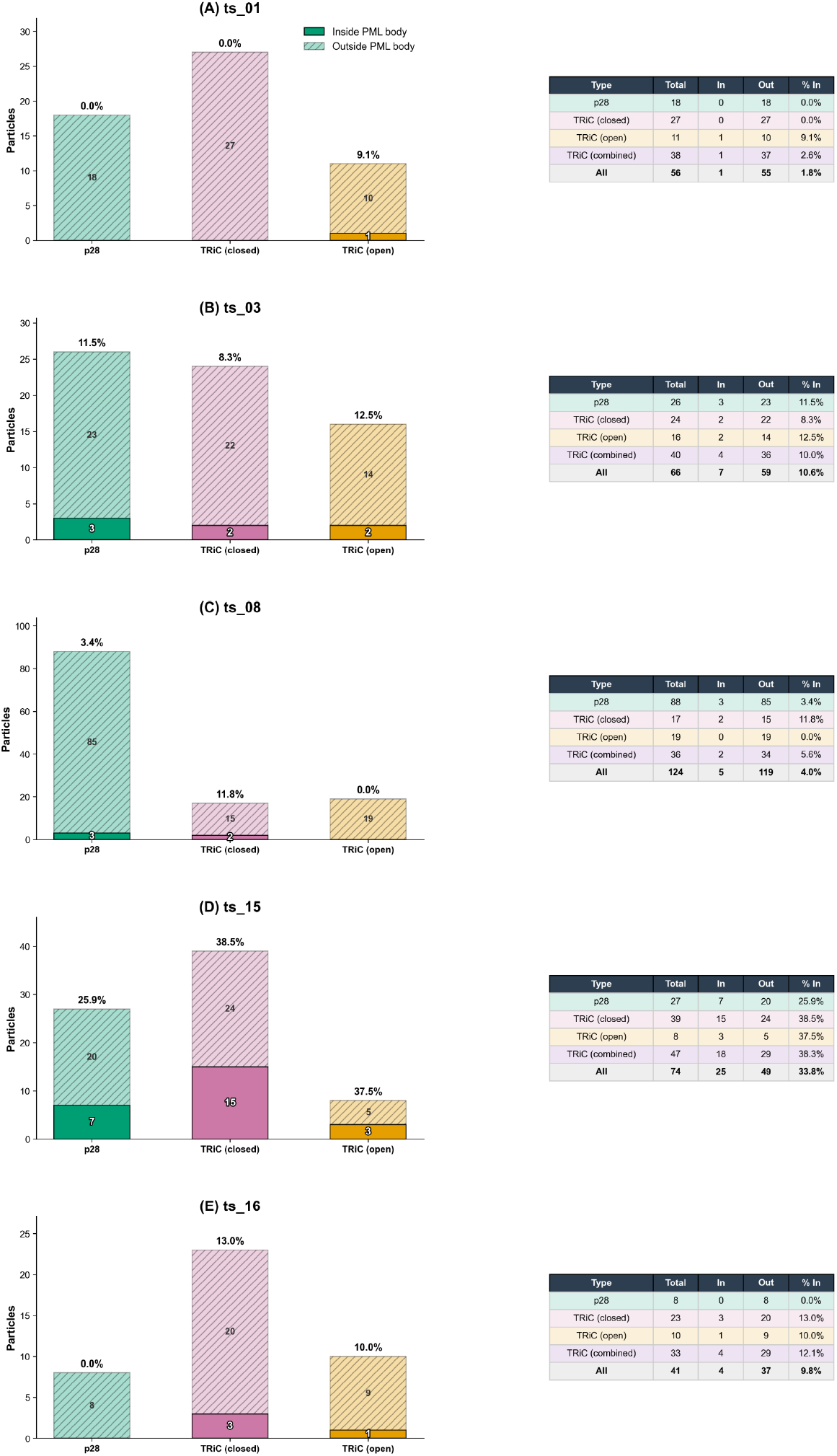
Quantification of PA28 and TRiC particle localisation within PML-I bodies. **(A-E)** Stacked bar charts and summary tables for each segmented tomogram (A, TS_01; B, TS_03; C, TS_08; D, TS_15; E, TS_16) showing the number of PA28, TRiC (closed), and TRiC (open) particles classified as inside (solid bars) or outside (hatched bars) the PML body. Percentages above each bar indicate the fraction of particles located within the PML body for each particle type. Summary tables list total, inside, and outside particle counts, with a combined TRiC row (closed + open) and an aggregate total.

**Supplementary Figure 11.**
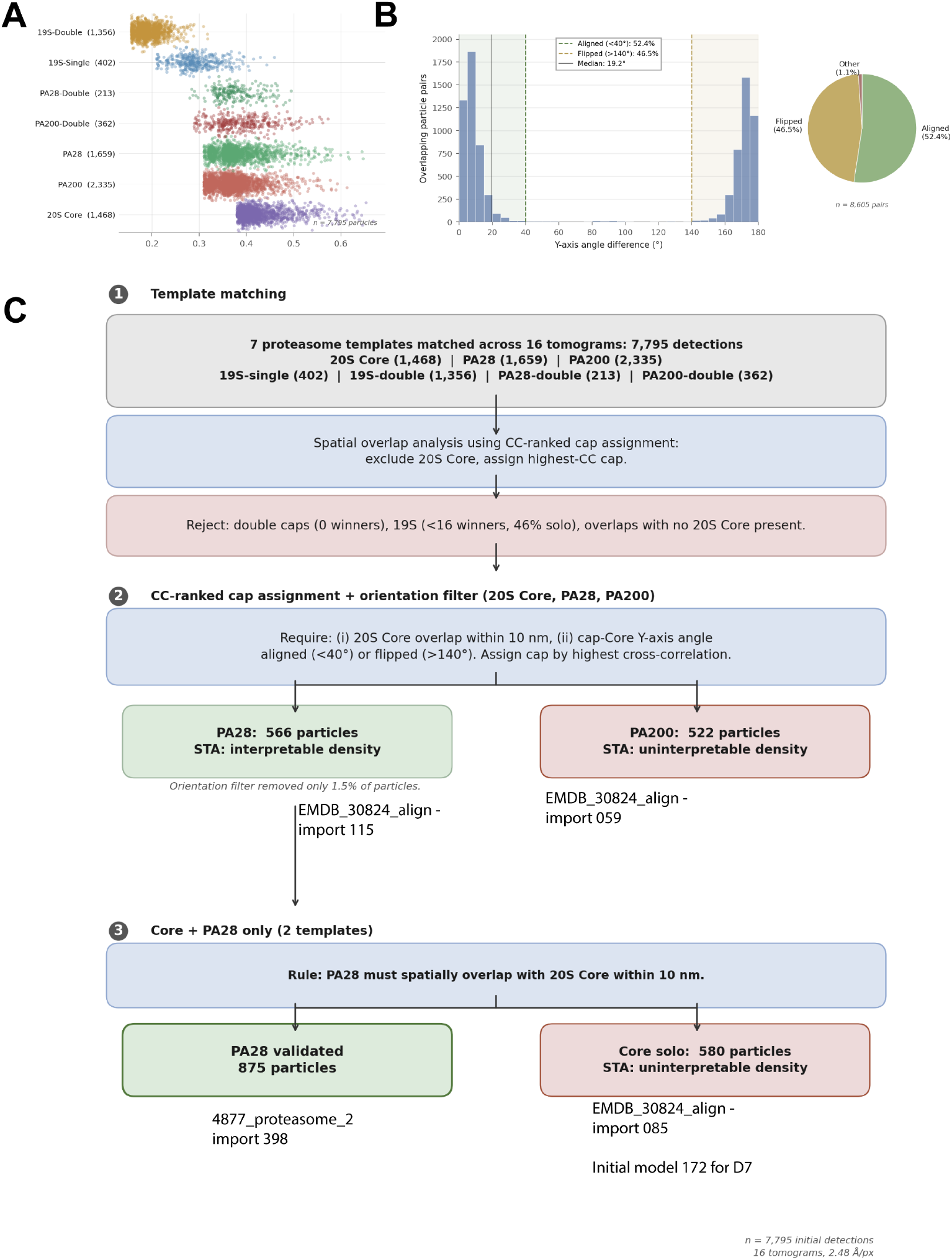
**(A)** Cross-correlation score distributions for seven proteasome-related templates across 16 cryo-tomograms (n = 7,795 initial detections). Templates are shown on the y-axis: 20S Core (1,468), PA200 (2,335), PA28 (1,659), PA200-Double (362), PA28-Double (213), 19S-Single (402), and 19S-Double (1,356). Y-axis angle difference between all inter-template overlapping detections within 10 nm (n = 8,605 pairs across all seven proteasome templates). The bimodal distribution shows two dominant populations: aligned (<40°, 52.4%) and flipped (>140°, 46.5%), consistent with equal-probability detection of the C2-symmetric 20S barrel in both orientations. Only 1.1% of pairs fall between 40-140° (pie chart, right). In Phase 2, each overlap group requires a Core anchor; orientation-validated PA28 and PA200 caps then compete by cross-correlation score, yielding 566 PA28 and 522 PA200 winners. **(C)** Schematic of the three-phase validation pipeline. Phase 1: seven proteasome templates matched across 16 tomograms yield 7,795 detections; spatial overlap analysis with CC-ranked cap assignment excludes 20S Core from competition; double-capped templates (0 winners) and 19S templates (<16 winners) are rejected. Phase 2: CC-ranked cap assignment with orientation filter applied to 20S Core, PA28, and PA200; requiring Core spatial overlap within 10 nm and cap-Core Y-axis angle aligned (<40 deg) or flipped (>140 deg) yields 566 PA28 particles (interpretable STA density) and 522 PA200 particles (uninterpretable STA density); orientation filter removed only 1.5% of particles. Phase 3: simplified two-template approach (Core + PA28 only), requiring PA28 to spatially overlap with 20S Core within 10 nm, yielding 875 validated PA28 particles and 580 core-solo particles (uninterpretable STA density).

**Supplementary Figure 12.**
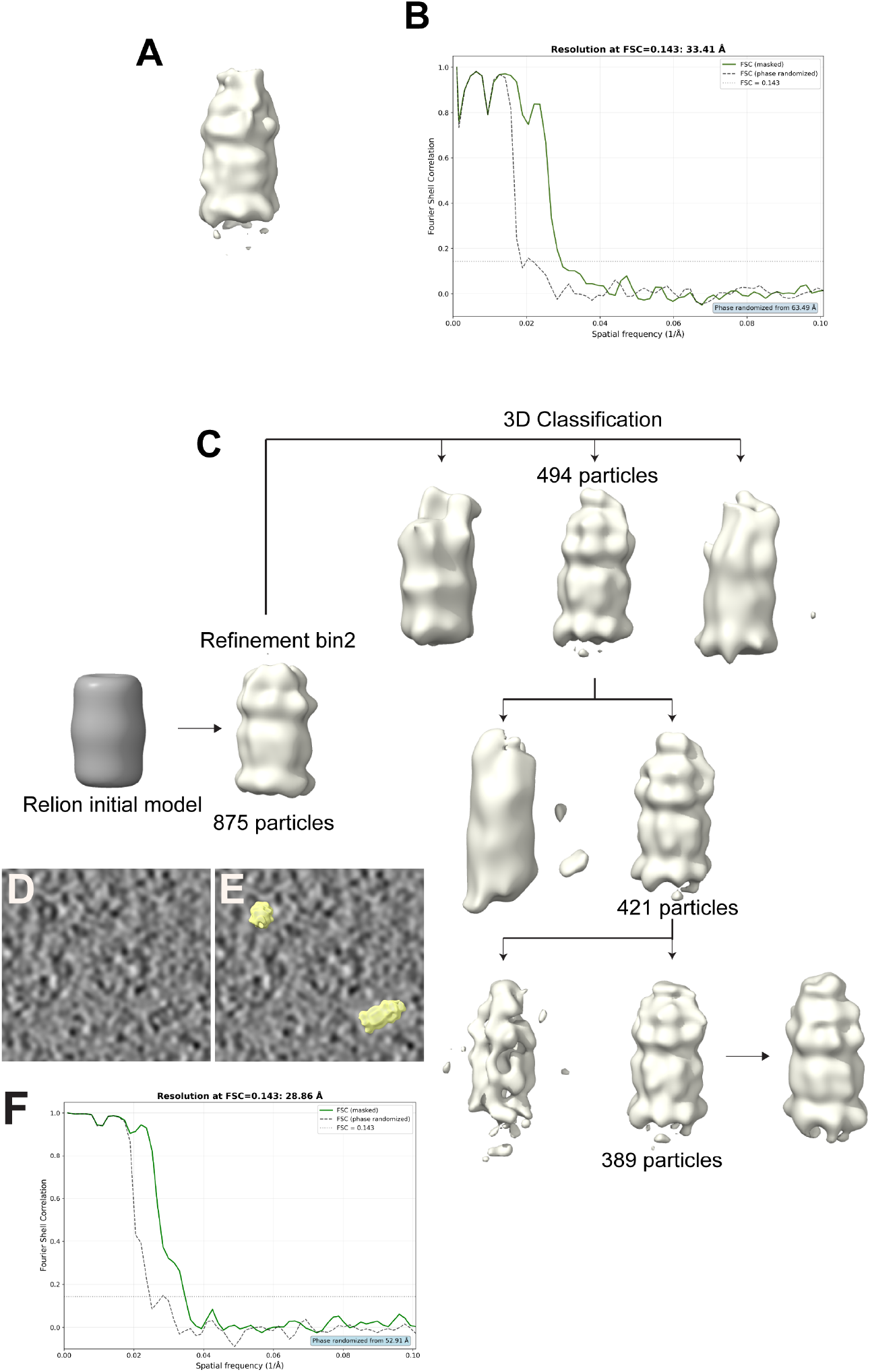
PA28 single-capped proteasome STA. **(A)** STA map of PA28 single-capped proteasomes derived from Phase 2 of the workflow shown in Sup. Fig. 11. **(B)** Fourier shell correlation (FSC) curve for the PA28 Phase 2 STA, showing a resolution of 33 Å at the 0.143 criterion. **(C)** STA pipeline for Phase 3 PA28 core-only overlaps. A de novo initial model was generated from 875 particles in RELION 4.0 and refined at bin2. Iterative 3D classification selected 494 particles, followed by further classification to 421 particles, and a final classification round yielding a best class of 389 particles. **(D)** Cryo-tomogram slice showing two PA28 single-capped proteasomes. **(E)** Same slice with PA28 particles highlighted in yellow. **(F)** FSC curve for the PA28 single-capped proteasome STA, yielding a resolution of 29 Å at FSC=0.143.

**Supplementary Figure 13.**
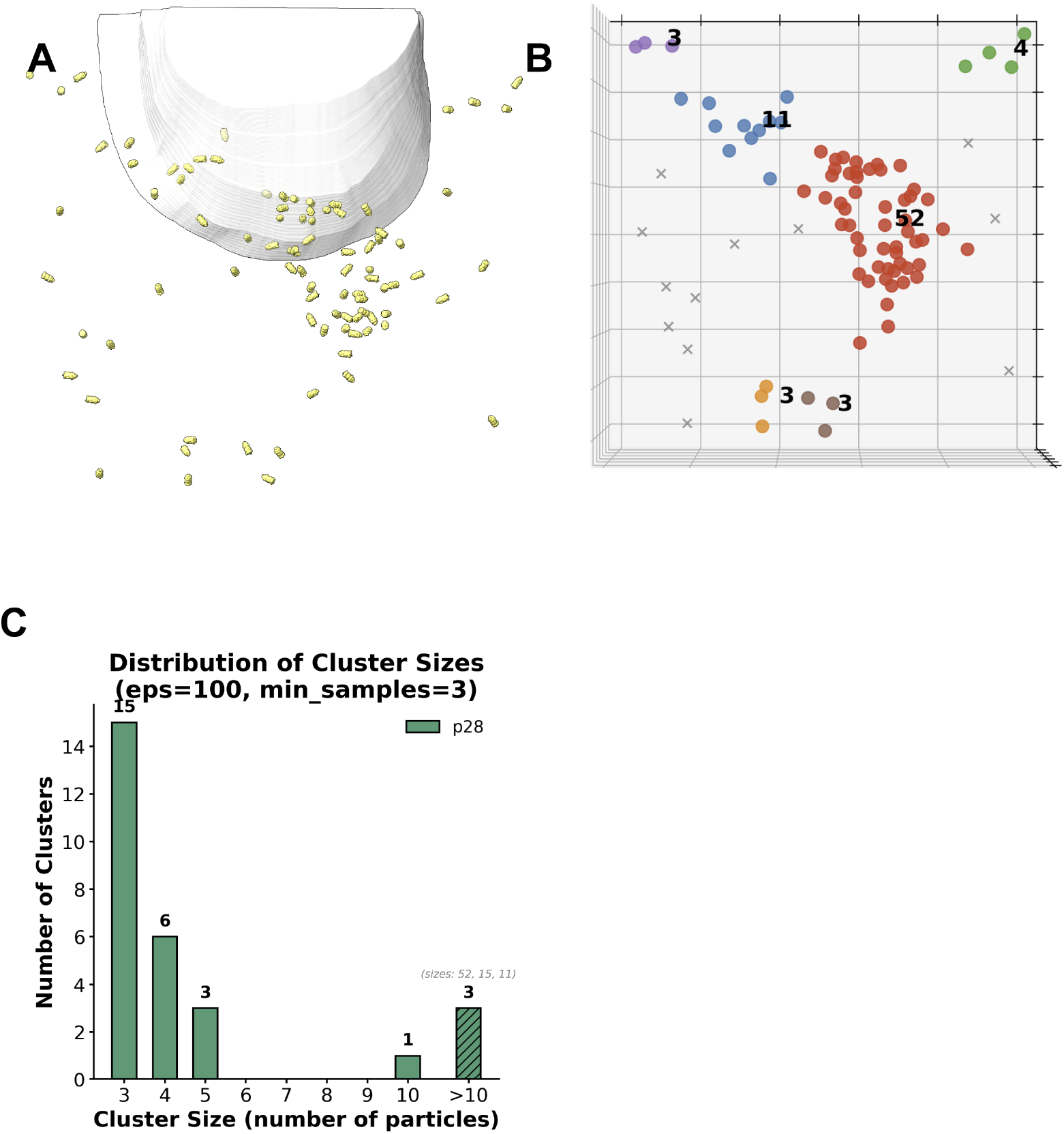
PA28 single-capped proteasome spatial distribution relative to the PML-I body. **(A)** PA28 single-capped proteasome STAs (yellow) mapped onto the Amira-generated PML-I body mesh surface (white), showing the spatial distribution of PA28 particles relative to the PML-I body. **(B)** DBSCAN clustering of PA28 particles in three dimensions, with cluster sizes indicated. **(C)** Histogram showing the distribution of DBSCAN cluster sizes for PA28 particles (eps = 100 bin4 pixels (∼99 nm), min_samples = 3). The majority of clusters contain 3–5 particles, with three larger clusters exceeding 10 particles (sizes: 52, 15, 11), indicating that PA28 particles form predominantly small, spatially dispersed clusters.

**Supplementary Figure 14.**
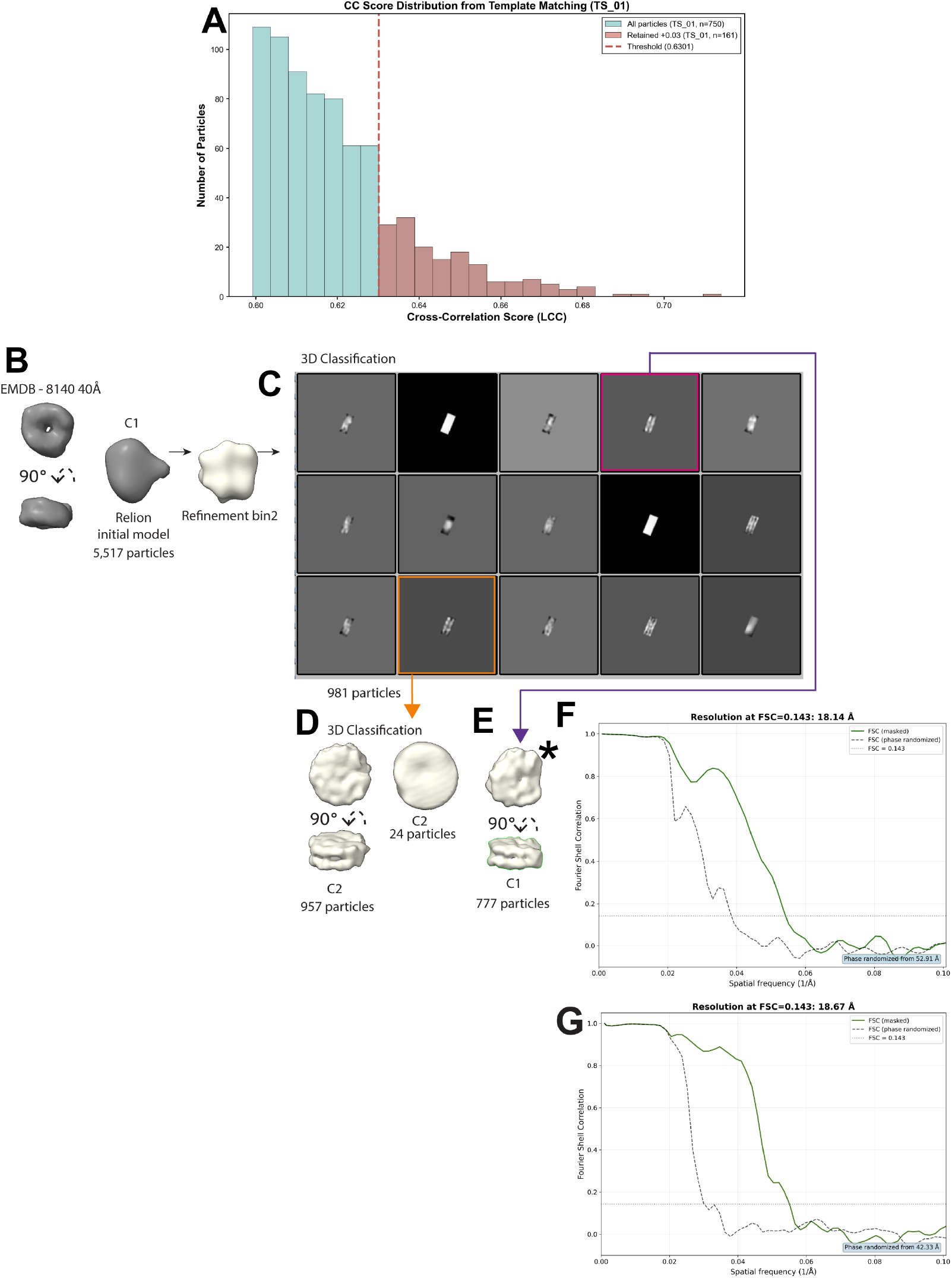
Nucleosome template matching and STA workflow. **(A)** Histogram of local cross-correlation (LCC) scores for all extracted particles (teal, n=750) and retained particles (pink, n=161) from tomogram TS_01. The per-tomogram threshold (dashed red line, LCC=0.6301) was set at +0.03 above the mean LCC score. Applying this threshold across all cryo-tomograms yielded 5,517 particles from an initial set of 22,680. **(B)** STA workflow showing the EMDB-8140 template (40 Å low-pass filtered, two orientations), RELION 4.0 initial model generation from 5,517 particles in C1 symmetry, and bin2 refinement. **(C)** First round of 3D classification into 15 classes; the orange-boxed class was selected for a second round of 3D classification and the pink-boxed class contained an extra density. **(D)** Second round of 3D classification in C2 symmetry yielding 957 particles, corresponding to the 19 Å nucleosome reconstruction. **(E)** C1 class of 777 particles; asterisk (*) indicates extra density adjacent to the nucleosome. **(F)** FSC curve for the nucleosome-with-extra-density STA map (777 particles), indicating a resolution of 18 Å at the FSC=0.143 criterion; phases randomised beyond 53 Å. **(G)** Fourier shell correlation (FSC) curve for the nucleosome STA, showing a resolution of 19 Å at the 0.143 criterion.

**Supplementary Figure 15.**
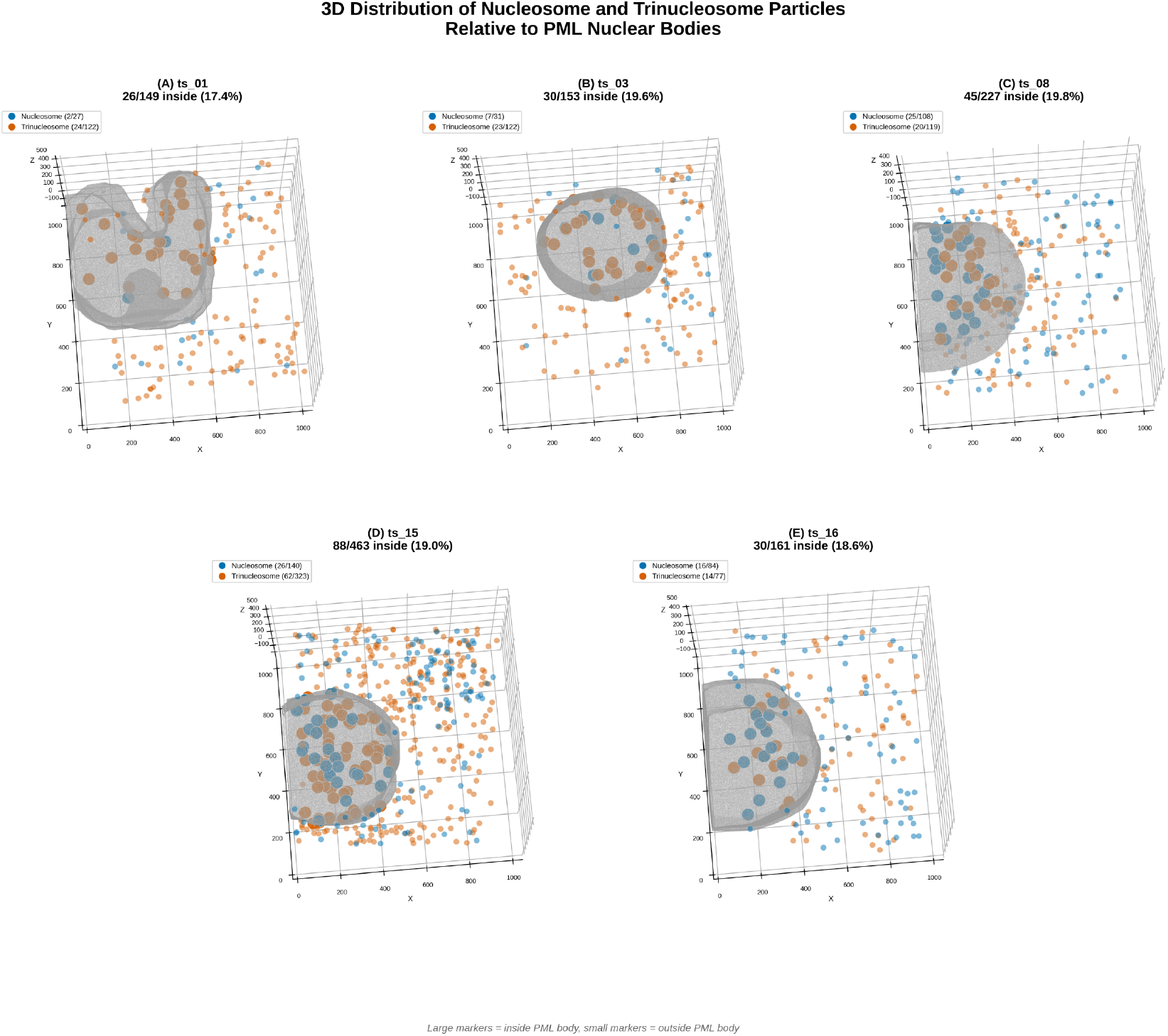
3D distribution of nucleosome and trinucleosome particles relative to PML nuclear bodies. **(A–E)** 3D renderings of PML body segmentations from five tomograms (A, TS_01; B, TS_03; C, TS_08; D, TS_15; E, TS_16), showing the spatial distribution of nucleosome (blue) and trinucleosome (vermillion) particles. PML body surface meshes (grey, semi-transparent) were segmented in Amira and rendered using isosurface extraction via marching cubes. Particle positions are from the refined STA particle sets and are displayed as spheres; large markers indicate particles classified as inside the PML body, small markers indicate particles outside.

**Supplementary Figure 16.**
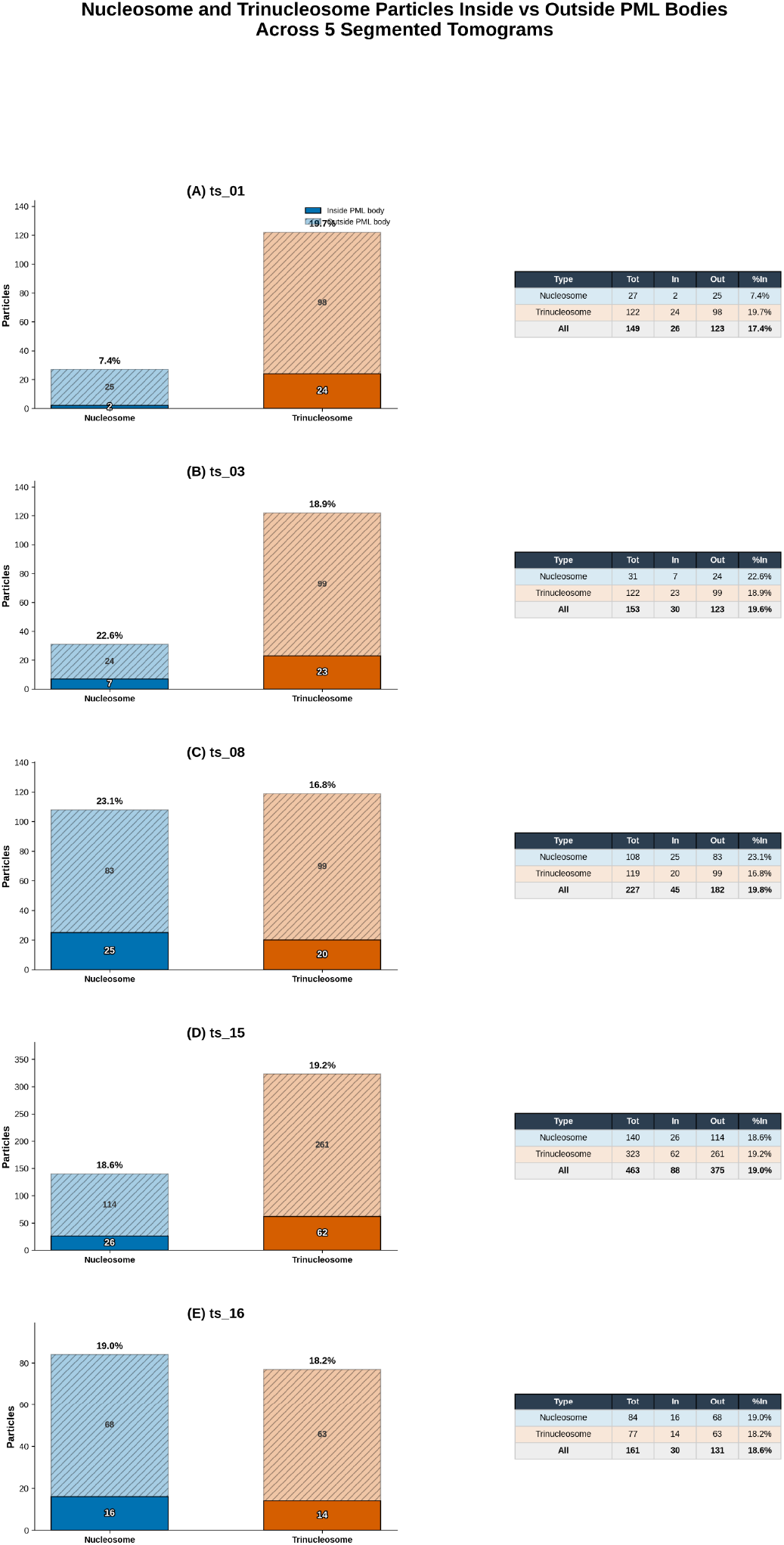
Quantification of nucleosome and trinucleosome particle localisation within PML nuclear bodies. **(A-E)** Stacked bar charts and summary tables for each segmented tomogram (A, TS_01; B, TS_03; C, TS_08; D, TS_15; E, TS_16) showing the number of nucleosome and trinucleosome particles classified as inside (solid bars) or outside (hatched bars) the PML body. Percentages above each bar indicate the fraction of particles located within the PML body for each particle type.

**Supplementary Figure 17.**
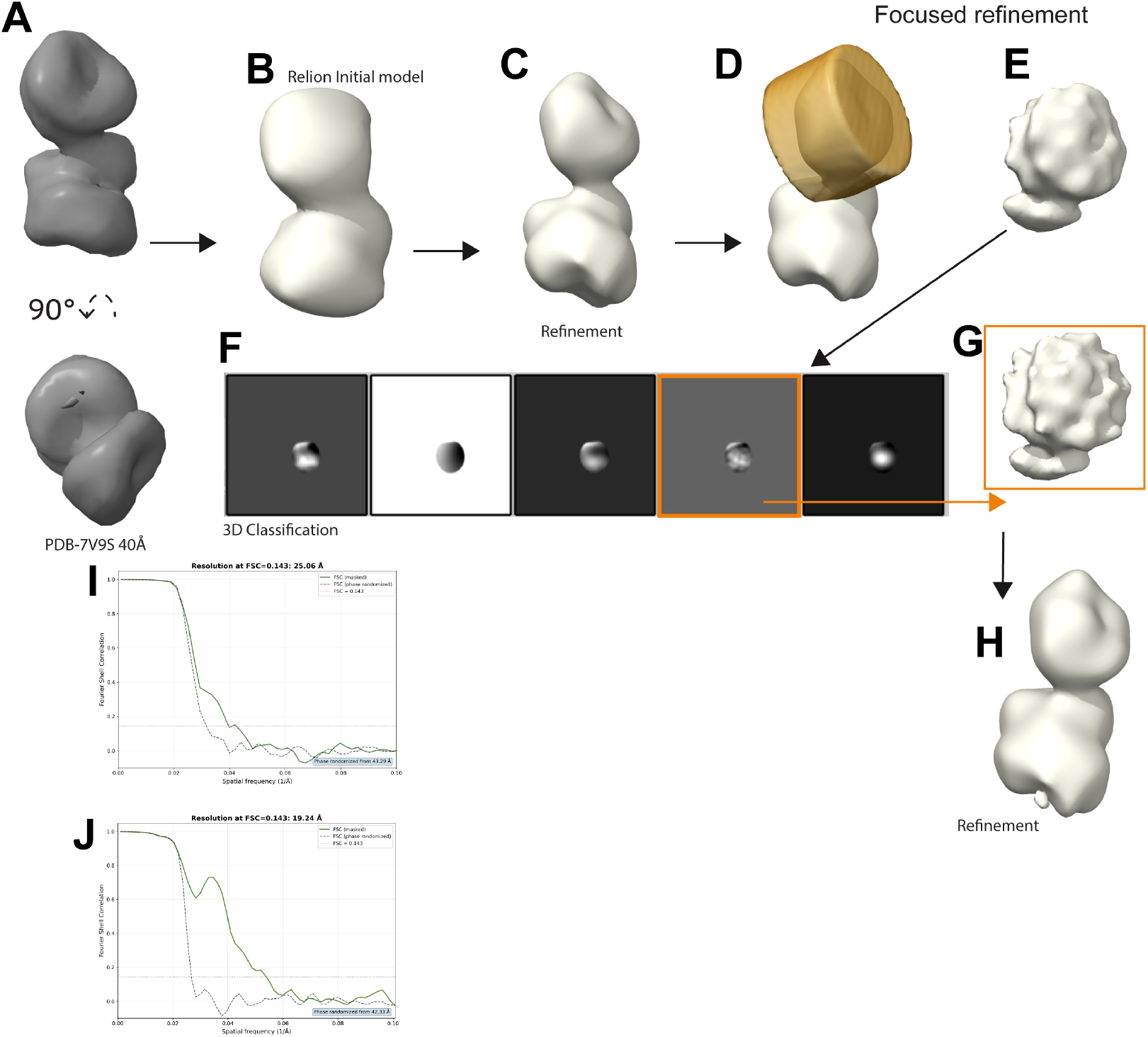
Open trinucleosome STA workflow. **(A)** Two views (90° apart) of the open trinucleosome template derived from PDB-7V9S, low-pass filtered to 40 Å. **(B)** De novo initial model generated in RELION. **(C)** Map following iterative 3D refinement. **(D)** Illustration of the focused refinement mask applied to the top nucleosome (orange). **(E)** Map following focused refinement. **(F)** 3D classification; the selected class is highlighted in orange. **(G)** Enlarged view of the selected class. **(H)** Final map following unrestricted refinement of the entire trinucleosome complex (2,287 particles), recovering the full complex after confirming that the top nucleosome signal exceeded the resolution of the original template. **(I)** Fourier shell correlation (FSC) curve for the initial trinucleosome refinement, showing a resolution of 25 Å at the 0.143 criterion. **(J)** Fourier shell correlation (FSC) curve for the focused refinement of the top nucleosome, showing a resolution of 19 Å at the 0.143 criterion.

**Supplementary Figure 18.**
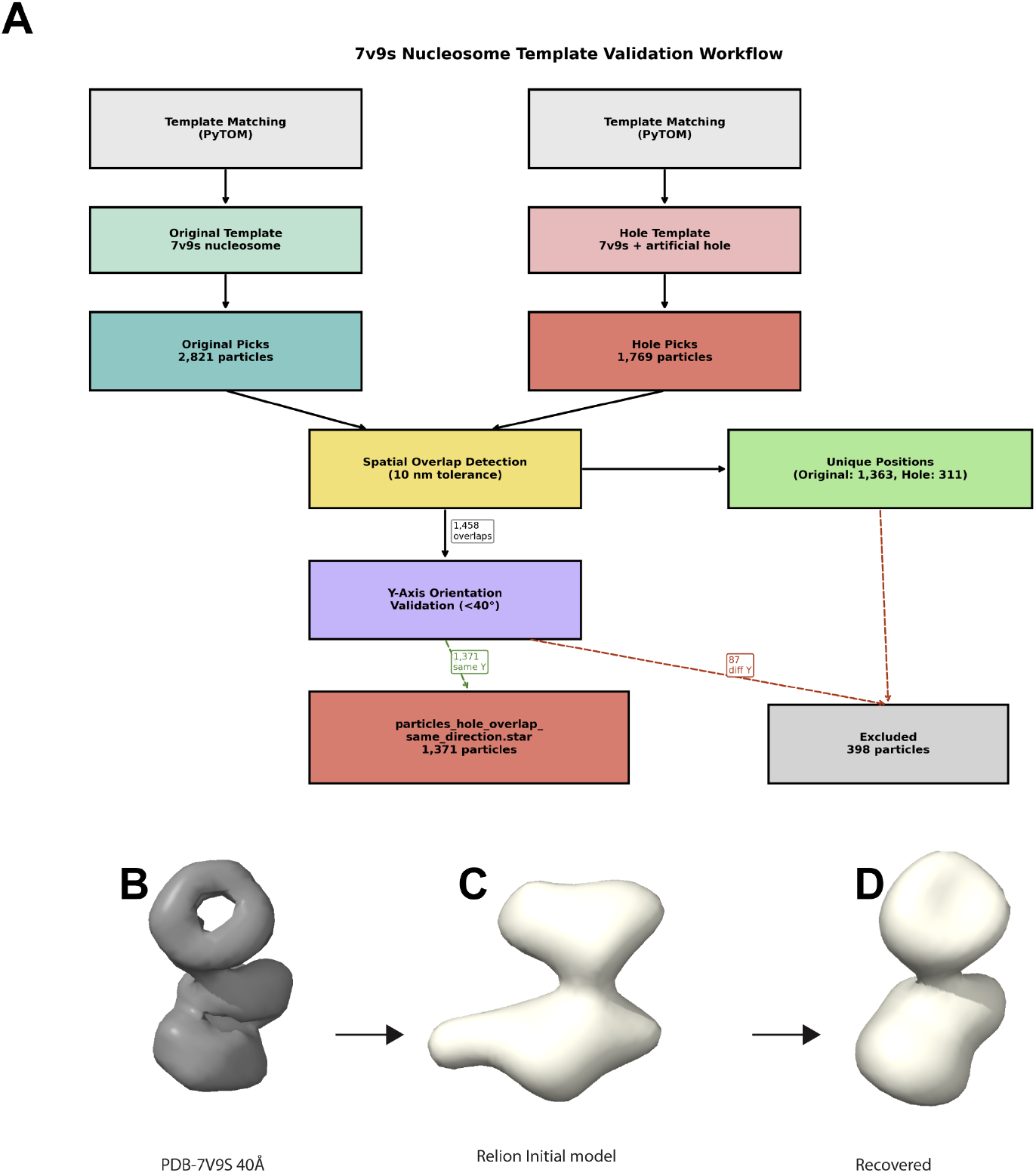
Trinucleosome template validation workflow. **(A)** Schematic of the template validation pipeline. Template matching was performed in PyTOM using both the original 7V9S trinucleosome template (2,821 particles) and a modified template with an artificial hole introduced into the top nucleosome (hole template; 1,769 particles). Spatial overlap detection with a 10 nm tolerance identified 1,458 overlapping positions; unique positions not detected by both templates were excluded (original. 1,363, hole: 311). Y-axis orientation validation (<40°) retained 1,371 overlapping particles in the same orientation, with 87 particles excluded. **(B)** PDB-7V9S filtered to 40 Å, used as the original trinucleosome template. **(C)** De novo initial model generated in RELION from the validated particle set. **(D)** Recovered map following STA, demonstrating that the missing density in the top nucleosome is restored, confirming that the trinucleosome detections represent genuine signal rather than template bias.

**Supplementary Figure 19.**
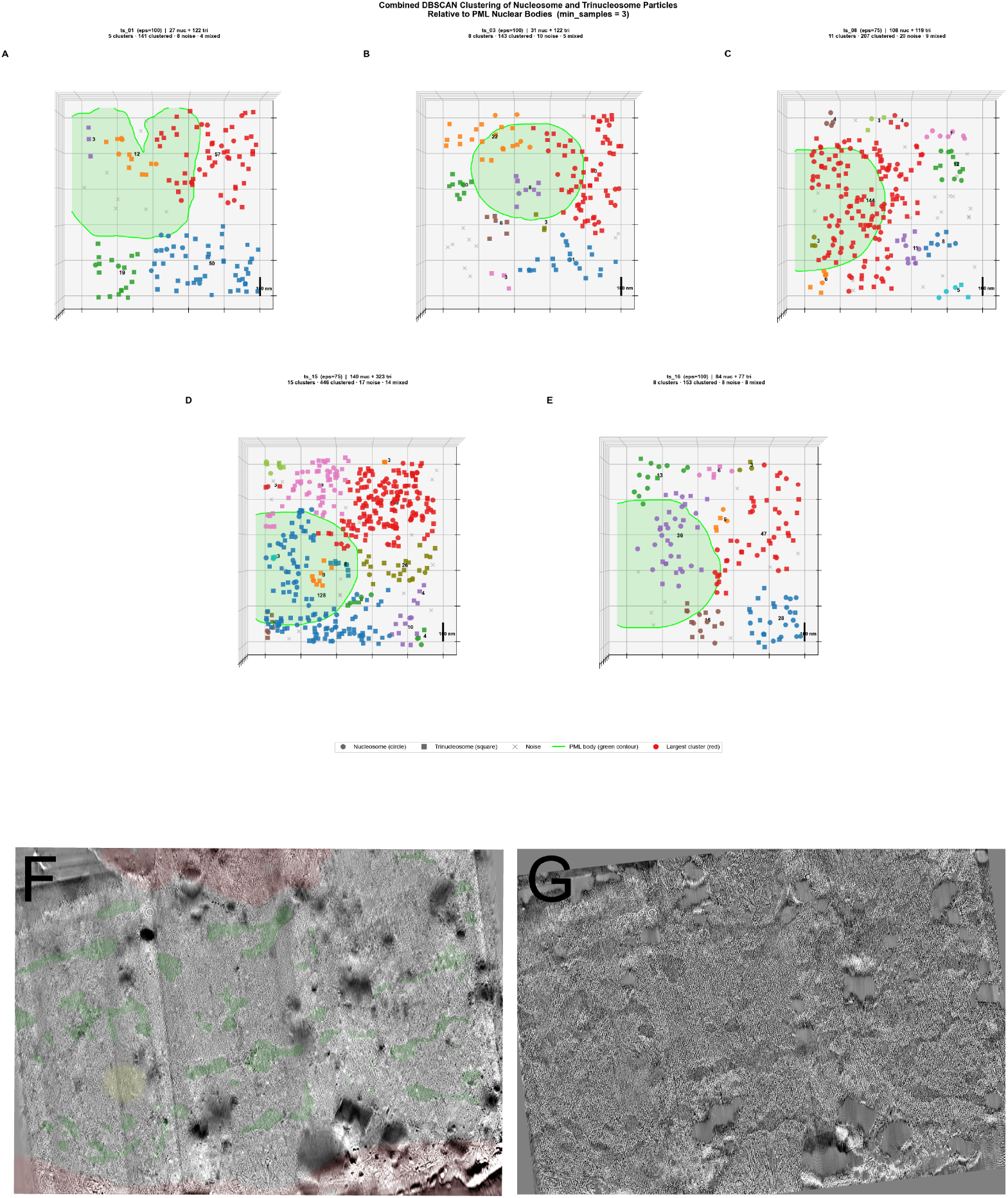
Combined DBSCAN spatial clustering of nucleosome and trinucleosome particles relative to PML nuclear bodies. **(A–E)** Top-down projections of DBSCAN clustering results for five segmented tomograms (A, TS_01; B, TS_03; C, TS_08; D, TS_15; E, TS_16). Nucleosome particles are shown as circles; trinucleosome particles are shown as squares. Each colour represents a distinct cluster; the largest cluster per tomogram is shown in red. Cluster particle counts are indicated at each cluster centroid. Grey crosses denote unclustered noise particles. The green filled region and outline indicate the maximum-intensity projection of the PML body segmentation onto the XY plane. DBSCAN parameters: eps = 100 bin4 pixels (∼99 nm) for TS_01, TS_03 and TS_16; eps = 75 bin4 pixels (∼74 nm) for TS_08 and TS_15; min_samples = 3. **(F, G)** Low-magnification cryo-tomograms of representative lamellae corresponding to the PML-I super-resolution puncta shown in Fig. 1. Overlay colours indicate cellular compartment identity: green, dense chromatin regions (nucleoplasm); red, cytoplasm.

**Supplementary Figure 20.**
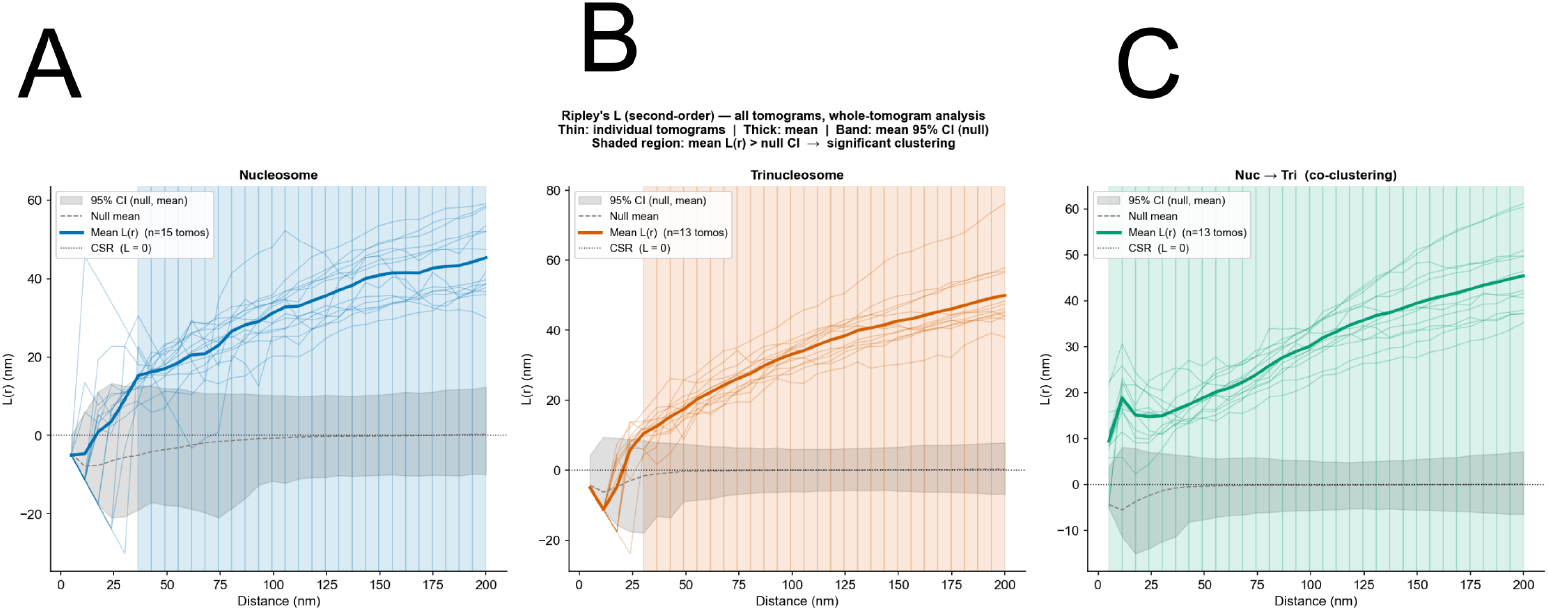
Ripley’s L function analysis of nucleosome and trinucleosome spatial clustering. Second-order Ripley’s L(r) for **(A)** nucleosomes (n = 15 tomograms), **(B)** trinucleosomes (n = 13 tomograms), and **(C)** bivariate nucleosome–trinucleosome co-clustering (n = 13 tomograms). Thick lines show the mean L(r); thin lines show individual tomograms. Grey band: 95% confidence interval of the null model (complete spatial randomness, CSR; 200 Monte Carlo simulations per tomogram). Coloured shading marks spatial scales where mean L(r) exceeds the null 95% CI (p < 0.05). Significant clustering was detected in every tomogram: 15/15 **(A)**, 13/13 **(B, C)**.

## Materials and Methods

### Cell culture

Human foreskin fibroblasts (HFFs) stably expressing eYFP-PML-I under doxycycline-inducible control were cultured as previously described (29, 30). Cells were initially maintained with 5 μg/ml hygromycin (Invitrogen, 10687-010) and 1 μg/ml puromycin (Sigma-Aldrich, P8833) for hTERT expression and selection, respectively. During experiments, HFFs were cultured in Minimum Essential Medium Eagle (MEM; Sigma-Aldrich M5650) supplemented with 10% fetal bovine serum, 2 mM L-glutamine (Life Technologies, 25030-024), 1 mM sodium pyruvate (Life Technologies, 11360-039), and 100 U/ml penicillin/streptomycin.

### 3D-SIM

Slides were coated with human fibronectin (20 mg/L) prepared in PBS (-Ca, -Mg). PML cells were grown in 8-well chamber slides and incubated with doxycycline for 7 hours. Cells were then fixed with 4% formaldehyde (diluted in PBS) for 15 minutes and washed twice in PBS, which was then left on. Samples in chamber slides were imaged using a ZEISS Elyra 7 microscope using Zen Black 3.0 SR FP2 software (version 16.0.21.306) with a 63×/1.4 NA oil immersion objective. Images were captured in lattice SIM mode, with a 27.5 µm grating and 13 phases, onto a PCO Edge 4.2 sCMOS camera. Z-stacks were captured at intervals of 110 nm and a 488 nm laser was used at 5% power. Processing was performed using SIM2 in 3D mode with input SNR medium, 5 iterations and regularisation 0, as per Rousset et al. (20).

### Immunofluorescence

HFF eYFP-PML-I cells were seeded onto coverslips at 1×10^5^ cells per coverslip and induced with 0.1 µg/ml doxycycline for 7 hours. Cells were washed in CSK buffer (10 mM HEPES, 100 mM NaCl, 300 mM sucrose, 3 mM MgCl_2_, 5 mM EGTA) and fixed/permeabilised in 1.8% formaldehyde and 0.5% Triton X-100 (Sigma-Aldrich, T-9284) for 10 min at RT. Following two PBS washes, coverslips were blocked for 30 min at RT in 0.45 µm-filtered PBS containing 2% FBS. Primary antibodies were applied individually in separate experiments: anti-Daxx (Upstate, 07-471; 1/50 dilution) or anti-MVP (Sigma-Aldrich, HPA002321; 1/500 dilution) (50 µl per coverslip, inverted staining). Coverslips were washed three times in PBS (5 min each) and incubated with goat anti-rabbit Alexa 555 secondary antibody (Invitrogen, A-21429; 1/1000 dilution). Following three PBS washes, three water washes, and air drying, coverslips were mounted onto glass slides with Citiflour AF1 (Agar Scientific, R1320). Positive controls (Daxx) and negative controls (secondary antibody only) were included in all experiments. Cell nuclei were counterstained with DAPI (1/1000 dilution; Sigma-Aldrich, D9542). Coverslips were examined using a Zeiss LSM 880 confocal microscope with a 63× Plan-Apochromat oil immersion objective (numerical aperture 1.4) using 405 nm, 488 nm, and 543 nm laser lines.

### Cryo-ET sample preparation

For cryo-ET sample preparation, Quantifoil SiO2 film R2/2 200-mesh gold grids were treated with laminin for 30-40 minutes and air-dried on Whatman 40 filter paper. Grids were placed in Mat-Tek glass-bottom dishes containing 2 ml culture media. HFFs were seeded onto grids and cultured for 12-14 hours to achieve optimal confluence for pFIB milling. eYFP-PML-I expression was induced by doxycycline treatment for 7 hours prior to vitrification. Vitrification was performed using a Vitrobot Mark IV (FEI). To ensure optimal structural preservation, excess medium was manually blotted from the grid backside using Whatman 40 filter paper 2 minutes before freezing. Subsequently, 3 μl of 8-9% glycerol in culture media was applied to the cell-bearing side as a cryoprotectant. Grids were immediately plunge-frozen in liquid ethane and stored in liquid nitrogen until pFIB milling.

### Arctis cryo-pFIB milling

For pFIB milling, frozen grids were clipped into autogrids and loaded into the Rosalind Franklin Institute Arctis cryo-pFIB/SEM (Thermo Fisher Scientific). SEM grid montages were collected using MAPS software to assess cell distribution and ice quality. Grids with optimal cell confluence, ice thickness, and structural integrity were selected for further pFIB milling.

### Cryo-fluorescence-guided pFIB target selection

Selected grids were rotated 180° on the stage to enable cryofluorescence imaging so the cells were facing the objective. Each grid square was systematically screened to identify cells exhibiting eYFP-PML-I nuclear bodies. The centre positions of nuclei containing eYFP-PML-I bodies were marked in the fluorescence images and their corresponding x,y coordinates were registered in the SEM montage using MAPS. Both green (eYFP-PML-I) and red channels were acquired with 500 ms exposure times to distinguish specific fluorescence from potential autofluorescence sources (60). Typically, 20 cells with bright eYFP-PML-I bodies were identified in the central region of each grid to ensure sufficient targets for successful lamella preparation.

### Lamellae preparation

Following fluorescence screening, the stage was rotated 180° back to its original position, placing cells facing the SEM and pFIB columns. A protective platinum layer was applied through a three-step process: (i) sputter coating with platinum for 2 minutes, (ii) organometallic platinum deposition using the gas injection system (GIS), and (iii) a second sputter coating cycle. This multi-layer coating strategy provided enhanced sample protection and charge dissipation during milling.

For lamella milling, MAPS software guided the stage to fluorescence-identified regions of interest. SEM images typically revealed oval-shaped protrusions at cell centres, which we interpreted as nuclei containing eYFP-PML-I bodies (Figure 1B). To achieve coincident SEM and FIB imaging planes, the stage Zheight was adjusted while monitoring FIB imaging, using grid features and cell boundaries as fiducial markers. Lamella positions were determined based on cellular topology, using features identifiable in the SEM x-y targeting coordinates that could also be visualised in the FIB view, while avoiding grid bars and thick ice accumulations. Lamella preparation was performed using the Thermo Fisher Arctis cryo-pFIB/SEM. Milling utilized both Argon and Xenon plasma ion beams operating at 30 kV accelerating voltage throughout all stages. A systematic multi-stage milling approach was employed with progressive current reduction to minimize beam-induced damage: typically, rough milling at 2 nA to remove bulk material, medium milling at 740 pA, fine milling at 200 pA, and sequential polishing steps at 60 pA and 20 pA. This protocol targeted a final lamella thickness of 150 to 200 nm and produced high-quality vitreous surfaces suitable for cryo-electron tomography. Data were collected across three cryo-ET sessions using Xe plasma FIB at 33,000× magnification (session 1), Xe pFIB at 53,000× (session 2), and Ar pFIB at 53,000× (session 3), yielding 13 vitreous lamellae in total.

### Post milling cryo-fluorescence screening of lamellae

After cryo-pFIB milling completion, each lamella was imaged in both green (eYFP-PML-I) and red channels to confirm retention of fluorescent signal and verify successful targeting of eYFP-PML-I bodies within the milled region. The platinum layer deposited during pFIB milling exhibited strong autofluorescence under cryo-conditions, which together with the lamella edges served as fiducial landmarks for correlating post-milling cryo-fluorescence images with TEM search maps, enabling precise tilt-series targeting. Approximately 40% of lamellae retained detectable eYFP-PML-I signal after milling.

### Cryo-SR-CLEM

Cryo-SR-CLEM was performed on cryo-lamellae using the VULCROM microscope as described in Falckenhayn et al. (34). A laser intensity of 200 W/cm^2^ of the 488 nm laser was used for fluorescence excitation and stimulation of photo-blinking. Data were recorded over the course of 2.6 h and binned to 500 ms per frame. Cryo-SMLM reconstructions were done as described in Kaufmann et al. (61). Single eYFP-PML-I molecules were localised with an average precision of 7.7 nm (Fig. 1U). Transformation of coordinate systems for correlation of cryo-SMLM with cryo-ET was done as described in Moser, PraŽák et al. (62). Localisations were overlaid onto cryo-ET tomogram slices to confirm that the eYFP-PML-I shell resides at the periphery of the finely textured body region (Fig. 1V–W).

### Tilt-series collection

Tomographic data were acquired using a Titan Krios G4 transmission electron microscope (Thermo Fisher Scientific) operated at 300 kV, Selectris energy filter, and Falcon 4i direct electron detector. An energy slit width of 10 eV was applied for all acquisitions. Dose-symmetric tilt series were collected using TOMO5 software (v5.17.0.6390, Thermo Fisher Scientific) with an automated image-shift/beam-shift acquisition strategy to maximize throughput. Data were recorded as movies in electron event representation (EER) format. Tilt series were then recorded of eYFPPML-I body territories in electron counting mode at a magnification of 33,000× and 53,000×. This corresponds to a pixel size of 3.770 Å and 2.48 Å, respectively, at the specimen level. Typically, the tilt scheme ranged from +60° to -60° in 2° increments. A -12° or -10° offset was applied to account for the lamella pre-tilt. A cumulative electron dose of 100-120 e/Å^2^ was distributed across the tilt series using a dose-symmetric acquisition scheme. Target defocus values ranged from -4 to -7 μm.

### Processing of tilt-series and reconstruction of cryotomograms

Motion correction and gain reference application for frame movies were performed using MotionCor2 (63). Tilt-series alignment and weighted back-projection tomographic reconstruction were carried out using AreTomo2 (64). Unbinned alignment parameters were exported in IMOD format (.xf transformation files) using the AreTomo2 -OutImod flag. These alignment parameters were subsequently used to generate pseudo-tomograms in RELION 4.0 (65) for STA. Output tomograms were binned by 4 for subsequent template matching, resulting in a pixel size of 9.92 Å.

### High-Contrast Dense Region Analysis

To assess whether large macromolecular densities are spatially associated with PML bodies, the bottom 15% of pixel intensities were identified after Gaussian smoothing (sigma = 20 pixels), applied to every third Z-slice through the PML body Z-range. A vertical reconstruction artifact along the left edge of each tomogram (X < 100 pixels) was masked and excluded from the analysis. The fraction of pixels classified as dense was then computed separately inside and outside the manually segmented PML body, pooled across all analysed slices, and the interior dense fraction was compared with that of the surrounding nucleoplasm.

### PML Body Network Segmentation and Analysis

Reconstructed cryo-electron tomograms were used at binned4 pixel size of 9.92 Angstrom. For each tomogram, a Z-slab encompassing the PML body was selected (TS_01: Z=93-255, TS_03: Z=82-245, TS_08: Z=85-249). Individual slices were independently z-scored, clipped to the +/-3 standard deviation range, and linearly rescaled to [0, 1] to standardise intensity prior to segmentation. **Denoising and Bright-Region Masking**. Each slice was denoised using total variation (TV) Chambolle denoising (weight = 0.05) to suppress high-frequency noise while preserving edges. To exclude low-contrast density regions, a large-scale background map was computed by Gaussian smoothing (sigma = 15 pixels) of the denoised image. Regions above the 75th percentile of this background map were removed from subsequent analysis. **Network Segmentation**. Contrast-limited adaptive histogram equalization (CLAHE; clip limit = 0.02, kernel size = 64 pixels) was applied to the denoised and background-subtracted image to enhance local contrast. A black top-hat transform (disk radius = 6 pixels) was used to extract small dark features corresponding to the fibrous density network. The top-hat response was normalised and thresholded at the 80th percentile of unmasked pixels to produce a binary segmentation. Small objects (<10 pixels) were removed, nearby fragments were connected by dilation (disk radius = 2 pixels), and small holes (<50 pixels) were filled to produce a clean binary network mask. **Skeletonization and Branch Point Detection**. The binary network mask was reduced to a one-pixel-wide skeleton using the Zhang-Suen thinning algorithm. Branch points were identified as skeleton pixels with three or more neighbours. **Grid-Based Connectivity Analysis**. Each tomogram slice was divided into a 12 × 12 grid. For each grid cell, skeleton fibre density and branch point density were computed and averaged across all Z-slices. A composite connectivity score was calculated from normalised fibre density (60% weight) and branch point density (40% weight). **PML Body Segmentation and Correlation**. PML bodies were manually segmented from denoised tomograms in Amira (described below). The PML body volume fraction per grid cell was computed over the Z-range containing the PML segmentation. Pearson correlation coefficients were calculated between PML volume fraction and connectivity score across all 144 grid cells per tomogram. **Software and Implementation**. Image processing was performed in Python 3.10 using scikit-image (v0.25.2), SciPy, and matplotlib. Analysis scripts are available at https://github.com/StephenC07/pml-body-network-analysis.

### Pore-Size Distribution Analysis

Pore diameters were measured per Z-slice on the 2D fibre skeleton produced above. For each non-skeleton pixel, the Euclidean distance to the nearest fibre was computed. Local maxima of this distance map mark the centres of the largest inscribed circles between fibres, and their values give the radii. Maxima were detected with a 5 by 5 maximum filter, accepted above 1 pixel, and rejected if within 5 pixels of the frame or if their inscribed circle extended beyond the PML interior mask. The mask was the maximum-Z projection of the manual PML body segmentation, used in 2D so that every Z-slice was evaluated over the same XY footprint. Each radius was multiplied by 2 × 0.992 nm (the bin-4 pixel size) to give a pore diameter. Diameters were pooled across all slices and the three cryo-tomograms (TS_01, TS_03, TS_08), yielding 363,437 measurements. The pooled mean is reported with standard deviation. Percentages of pores exceeding a nucleosome (∼11 nm), TRiC (∼17 nm) and PA28-capped 20S proteasome (∼20 nm) were calculated as integer fractions. Pore diameters reflect 2D cross-sectional gap sizes, not isotropic 3D voids. The measurement was validated on a synthetic Voronoi-mesh phantom of known inscribed-circle diameters, recovering the median to within 8 percent for pores of 15 nm and above, with a slight underestimate of up to about 14 percent for pores below 12 nm. This underestimate is conservative for the exclusion analysis, as it would only lower the fraction of pores scored above each complex diameter. Analysis and validation scripts are available at https://github.com/StephenC07/pml-body-network-analysis.

### Template Matching

Template matching was performed using PyTOM v0.7.1 on 4× binned cryo-tomograms (pixel size: 9.92 Å). Templates were generated from atomic models using a two-step process. First, PDB structures were converted to density maps at 25 Å resolution using the molmap command in ChimeraX (molmap #1 25 gridSpacing 9.92) with the binned 4 pixel size. Templates were then generated using PyTOM v0.3.1 pytom_create_template.py with the following parameters: –input-voxel 9.92 –output-voxel 9.92 –centre –ctfcorrection -z [defocus] -a 0.07 -v 300 –Cs 2.7 –flip-phase –invert -b 80 –low-pass 40. The CTF correction parameters corresponded to our microscope settings (spherical aberration Cs = 2.7 mm, amplitude contrast = 0.07, voltage = 300 kV) with tomogram-specific defocus values of 4, 5, 6, and 7 μm.

Templates derived from EMDB density entries were generated using identical parameters minus the molmap step, except the input pixel size was adjusted to match the respective EMDB map pixel size. Spherical masks were generated using PyTOM v0.3 (pytom_create_mask.py -b 96 -o mask.mrc -r # –radius-minor1 # –radius-minor2 # -s 1 –voxel-size 9.92) and resampled using ChimeraX’s vop resample command (vop resample #1 onGrid #0).

### Template Matching Execution

Template matching was executed using pytom_match_template.py with the following parameters: -t [template_path] -v [tomogram_path] -m [mask_path] -d [results_directory] -a [tilt_angles_file] –angular-search 7.00 –voxel-size 9.92 –g 0 1 2 –search-z –search-x. The angular search step of 7.00° provided sufficient sampling while maintaining computational efficiency. The Z-search range was constrained to the lamella thickness to restrict template matching to the cellular volume and exclude false-positive matches from the lamella surfaces or surrounding ice. The X-search range was constrained to avoid reconstruction artefacts generated by AreTomo2. Specific PDB/EMDB accession numbers used in this study are detailed in Sup. Fig. 6. Correlation score maps generated by PyTOM were manually inspected using a custom napari viewer (simple_threshold_view_scale.py). This plugin enabled interactive adjustment of the cross-correlation threshold while simultaneously displaying the corresponding tomographic data. The viewer was used for inspection and threshold selection only and does not itself extract particles. The plugin also supported generation of movies for documentation, with an automated scale bar for spatial reference (available at https://github.com/StephenC07/pytom-cc-threshold-viewer). Candidate particles above the chosen threshold were extracted from the score maps using PyTOM (pytom_extract_candidates.py) and written as RELION 4.0 STAR files with x,y,z coordinates and Euler angles at the unbinned pixel size of 2.48 Å/pixel. STAR files from individual tomograms were subsequently merged into a single combined STAR file using pytom_merge_stars.py prior to import into RELION 4.0.

### STA using RELION 4.0 (65)

#### TRiC, closed conformation

335 particles were imported into RELION 4.0 and an initial model was generated de novo in D8 symmetry using 100 vdam mini-batches. Particles were refined at binning 4 in C1. The 3D reference map was reconstructed directly from 2D tilt series projections using the RELION tomo_reconstruct_particle and pseudo-tomograms were produced at bin 2. The reconstructed particle was used as a reference for bin 2 refinement. 3D classification was performed with 7 classes at T=0.5 to remove noise particles, yielding a single good class of 303 particles in C1 followed by a second round of 3D-classification at T=1 with 3 classes, yielding a final class of 297 particles. A final round of refinement was performed in D8, yielding a resolution of 22 Å at FSC = 0.143.

#### TRiC, open conformation

413 particles were imported into RELION 4.0 and an initial model was generated de novo in C1 symmetry using 50 vdam mini-batches. Particles were refined at binning 4 in C1, and the 3D reference map was reconstructed directly from 2D tilt series projections using the RELION tomo_reconstruct_particle, with pseudo-tomograms produced at bin 2. The reconstructed particle was used as a reference for bin 2 refinement. Iterative 3D classification was performed to remove noise particles: a first round with 3 classes at T=1, a second round with 3 classes at T=1, followed by 2 classes at T=1 and a final round with 2 classes at T=2, yielding 276 particles. A final round of refinement in C1 at binning 2 yielded a resolution of 32 Å at FSC = 0.143.

#### PA28, Phase 2

566 particles were imported into RELION 4.0 and an initial model was generated de novo in C1 symmetry using 50 vdam mini-batches. Particles were refined at bin2 in C1, and the 3D reference map was reconstructed directly from 2D tilt series projections and pseudo-tomograms generated at bin2. A second round of C1 refinement was performed at bin2. 3D classification (T = 2, 3 classes, no angular sampling) identified one interpretable class containing 538 particles. Two further rounds of refinement at bin2 in C1 yielded a final reconstruction at 33 Å resolution (FSC = 0.143 criterion; Sup. Fig. 12A, B).

#### PA28, Phase 3

875 particles were imported into RELION 4.0 and an initial model was generated de novo in C1 symmetry using 120 vdam mini-batches. Particles were refined at bin4 in C1. The 3D reference map was then reconstructed directly from 2D tilt series projections and pseudo-tomograms were generated at bin2, followed by two rounds of refinement in C1 prior to 3D classification. 3D classification (T = 2, 3 classes, no angular sampling) selected 494 particles; a second round (T = 2, 2 classes, no angular sampling) yielded 421 particles; and a final round (T = 2, 2 classes, no angular sampling) identified a final class of 389 particles. A final round of refinement yielded a reconstruction at 29 Å resolution (FSC = 0.143 criterion; Fig. 2Q–S; Sup. Fig. 12C–F).

#### Nucleosome

Template matching against all 16 tomograms using EMDB-8140 yielded 22,680 initial particle candidates (mean local cross-correlation, LCC = 0.519). To select high-confidence detections, a per-tomogram CC score threshold of mean LCC + 0.03 was applied (0.6301 for TS_01); particles from tomogram TS_162 were additionally excluded owing to the presence of embedded gold fiducials, retaining 5,517 particles (mean LCC = 0.548; Sup. Fig. 14A). These 5,517 particles were imported into RELION 4.0 and an initial model was generated de novo in C1 symmetry using 50 vdam mini-batches. Particles were refined at bin4 in C1. The 3D reference map was then reconstructed directly from 2D tilt-series projections and pseudo-tomograms were generated, followed by refinement at bin2 in C1. 3D classification with 15 classes (T = 2, with angular sampling; angular sampling interval 1.8°, local angular search range 15°) identified two classes of interest: 981 particles corresponding to a nucleosome-like density and 777 particles exhibiting extra density at one linker-nucleosome boundary (Sup. Fig. 14B, C). The 981-particle class was taken forward into a second round of 3D classification in C2 symmetry (2 classes, T = 1), yielding a final reconstruction of 957 particles. No further classification was performed on the 777-particle extra-density class. Final refinement yielded a 19 Å reconstruction from the 957-particle class and an 18 Å reconstruction from the 777-particle extra-density class (FSC = 0.143 criterion, Sup. Fig. 14D and F).

#### Trinucleosome

Template matching against all 16 tomograms using PDB-7V9S yielded 2,821 initial particle candidates; particles from tomograms TS_04 and TS_162 were subsequently excluded as CC score peaks in these cryo-tomograms correlated predominantly with noise. These particles were imported into RELION 4.0 and an initial model was generated de novo in C1 symmetry using 30 vdam mini-batches. Particles were refined at bin4 in C1. The 3D reference map was reconstructed directly from 2D tilt-series projections and pseudo-tomograms were generated, followed by a second round of refinement at unbinned pixel size (2.48 Å/pixel) in C1, yielding a reconstruction of the full trinucleosome at 25 Å resolution (FSC = 0.143 criterion). Focused classification was then performed using a cylindrical mask targeting a single nucleosome subunit, generated in ChimeraX (66) using a cylinder, converted to a binary mask, smoothed with a Gaussian filter (σ = 2), and resampled to match the map grid prior to use. Focused 3D classification (5 classes, T = 1, no angular sampling) identified one class containing 2,287 particles; focused refinement on the top nucleosome subunit at bin2 (4.96 Å/pixel) yielded a map at 19 Å resolution (FSC = 0.143 criterion). Final refinement was performed using a trinucleosome mask generated in RELION 4.0 to align the global structure. A total of 2,287 trinucleosome particles were used for spatial clustering analysis. Particles from cryo-tomogram TS_17 were filtered out through 3D-classification prior to clustering, leaving 13 tomograms.

#### Trinucleosome template matching and STA validation

To validate trinucleosome particles detected by template matching, we employed a dual-template strategy. Two templates were derived from the 7V9S trinucleosome structure: (2) the original intact structure, yielding 2,821 initial particle candidates, and (3) a modified template in which an artificial hole was introduced into one nucleosome subunit, yielding a further 1,769 detections. Template matching was performed independently with each template across all 16 tomograms at a pixel size of 9.92 Å. Detections from both templates were cross-referenced: 1,458 overlapping pairs localised within 10 nm of one another were identified. For each overlapping particle pair, orientation consistency was assessed by comparing Y-axis direction vectors derived from RELION ZYZ Euler angles. Of the 1,458 pairs, 1,371 exhibited consistent Y-axis orientations (mean angular deviation 8.9°) and were retained; those exceeding the 40° threshold were excluded. The final validated particle set retained only hole-template coordinates and orientations from orientation-consistent overlaps. STA of the hole-modified particles was performed independently: an initial model was generated de novo in C1 symmetry using 30 vdam mini-batches, followed by refinement at bin2 (4.96 Å/pixel) in C1. Template matching was additionally performed using alternative telomeric nucleosome array structures: a tetranucleosome (PDB-7V9K), and a tetranucleosome in an open state (PDB-7VA4); none yielded a subtomogram average exceeding the resolution obtained with PDB-7V9S.

#### PyTOM cross-template overlap and orientation analysis

Each template in this pipeline was first oriented manually in UCSF ChimeraX with its principal axis aligned to the reference Z axis: the TRiC open and closed templates along the chaperonin barrel axis, the seven 20S proteasome templates (20S core, PA28, PA200, 19S-single, 19S-double, PA28-double, PA200-double) along the proteasome barrel axis, and the original and hole-modified 7V9S trinucleosome templates along the stack axis. This common reference orientation ensured that the identical 7° angular sampling grid used by pytom-match-template explored the same orientation space for every template, so that Euler angles of detections from different templates were directly comparable. Template matched particles exported by pytom-match-template were written as STAR files retaining the PyTOM Euler angles in the _rl-nAngleRot, _rlnAngleTilt and _rlnAnglePsi columns. PyTOM’s proper-Euler ZYZ convention (source: pytom/angles/angleFnc.py, zyzToMat) defines the active reference-to-lab rotation as R = Rz(ψ)·Ry(θ)·Rz(φ), with φ = rot, θ = tilt, ψ = psi, and the lab-frame direction of the reference Y axis given by R·[0, 1, 0]. All PyTOM-output analyses described below use this matrix; the RELION-refined barrel-axis analysis of TRiC clusters (reported separately) uses RELION’s own matrix, which differs in composition order and must not be confused with the PyTOM expression above. For each pair of template STAR files, particles in the same tomogram were grouped into spatial clusters within a 10 nm radius (unbinned pixel size 2.48 Å). Within every overlap, orientation consistency between detections was assessed by computing the angle between their lab-frame Y axes.

For pairs compared against a C2-symmetric reference (20S proteasome core), overlaps were classified as aligned (Y-axis angle < 40°) or flipped (> 140°) and the remaining pairs excluded, reflecting the barrel’s C2 equivalence under 180° flips. For pairs compared against a non-symmetric reference (the original 7V9S trinucleosome template vs. a hole-modified variant of the same template), only the aligned criterion (< 40°) was applied. For TRiC (open vs closed conformations) and for proteasome cap assignment (Phase 2 and Phase 3), the template at each retained overlap was identified as the detection with the highest cross-correlation score (_rlnLCCmax). For the trinucleosome, where both templates represent the same structure, no cross-correlation competition was applied; hole-template detections that passed both the spatial overlap and the Y-axis consistency check were retained as the validated particle set.

#### Denoising of cryo-tomograms for presentation

Denoising was performed using IsoNet (31) and total variation (TV) denoising. IsoNet was applied using deconvolution parameters (Decon-vStrength = 1.5, SnrFalloff = 1.5). TV denoising was performed in Dragonfly (32) (Object Research Systems, Montreal, Canada).

#### Visual proteomic analysis for TRiC

Spatial clustering of TRiC particles was performed independently for each conformational class (closed and open) using DBSCAN (38) (Density-Based Spatial Clustering of Applications with Noise) as implemented in scikit-learn (67). Particle coordinates were extracted from per-tomogram RELION STAR files and clustering was performed in 3D coordinate space with parameters eps ≈ 37 nm and min_samples = 2. The eps value of ∼37 nm was chosen to capture extended clusters in which particles are connected through chains of neighbours; clusters containing two or more particles were retained and unclustered particles were classified as noise. To test whether clustering frequency differed with proximity to PML-I bodies, tomograms were classified as PML-body-proximal (n = 7) or distal (n = 9) according to whether an eYFP-PML-I body was captured, and the per-tomogram fraction of clustered closed-TRiC particles was compared between groups with a two-sided Mann-Whitney U test. To measure packing distances within clusters, nearest-neighbour distances were calculated for each particle. Only mutual nearest-neighbour pairs were recorded, where both particles identified each other as their closest neighbour, to avoid double counting. This mutual constraint prevents spurious long-distance pairings in clusters containing more than two particles. Centre-to-centre distances were computed in pixels and converted to nanometres (pixel size = 2.48 A/pixel). Only pairs within 20 nm were retained for distance and orientation analysis, consistent with the cluster threshold used by Xing et al. (39).

To determine the relative orientation of paired TRiC complexes, Euler angles for closed TRiC were taken from D8-symmetry-refined STA output and for open TRiC from C1 refinement (RELION). The barrel axis of each particle was defined as the third row of the RELION rotation matrix A, computed from Euler angles (rot, tilt, psi) following the RELION convention (65):

<preformat>barrel = [sin(tilt)*cos(rot), sin(tilt)*sin(rot), cos(tilt)]</preformat>

This corresponds to the reference Z-axis (the barrel axis) transformed into tomogram coordinates, consistent with the ChimeraX Z-axis alignment of the TRiC templates during template preparation described above. The angle between barrel axes of paired particles was computed as the arc-cosine of their dot product. Because TRiC possesses D8 point group symmetry, the equatorial C2 elements within D8 render the barrel axis direction ambiguous (the barrel viewed from either end is equivalent). Barrel axis angles were therefore folded to the 0-90 degree range by taking min(angle, 180 angle). The connection angles were likewise folded to the 0-90 degree range before the angular thresholds below were applied.

Pairs were classified into three contact geometries based on two angular measurements: (i) the relative barrel axis angle between paired particles, and (ii) the connection angle, defined as the angle between the inter-particle vector and each particle’s barrel axis. Equator-to-equator contacts were defined as pairs with aligned barrel axes (relative barrel axis angle < 40 degrees) and connection angles >= 40 degrees for both particles, indicating the inter-particle vector lies perpendicular to the barrel axes. Head-to-head contacts were defined as pairs with aligned barrel axes and connection angles < 20 degrees for both particles, indicating the inter-particle vector lies along the barrel axes. All remaining pairs were classified as head-to-equator. This group comprised pairs with near-perpendicular barrel axes (relative barrel axis angle >= 40 degrees) together with aligned-barrel pairs whose connection angles met neither the head-to-head nor the equator-to-equator criterion. These residual aligned-barrel pairs were reported separately as an intermediate class. Code implementing the DBSCAN-based TRiC pair-orientation classification is available at https://github.com/StephenC07/tric-cluster-orientation-analysis.

### DBSCAN clustering analysis for PA28 single-capped proteasome, nucleosomes and trinucleosomes

To assess the spatial organisation of nucleosome and PA28 particles within tomograms, density-based clustering was performed using DBSCAN (38) (Density-Based Spatial Clustering of Applications with Noise) as implemented in scikit-learn (67). Particle coordinates from template matching were scaled to bin4 pixel space (divided by 4) prior to clustering. DBSCAN was run with min_samples = 3 and a tomogram-specific eps: 100 bin4 pixels (∼99 nm) for TS_01, TS_03 and TS_16, and 75 bin4 pixels (∼74 nm) for TS_08 and TS_15, reflecting differences in particle density across tomograms; a smaller eps was applied to TS_08 and TS_15 to avoid over-merging spatially distinct clusters in these more densely populated datasets. Particles not assigned to any cluster were classified as noise. For each tomogram, clusters were visualised as 3D scatter plots with individual clusters colour-coded and particle counts annotated at cluster centroids.

To formally assess deviation from complete spatial randomness (CSR), the Ripley’s L function was computed for nucleosome and trinucleosome particles across 15 and 13 tomograms respectively. The edge-corrected Ripley’s K estimator (Ohser–Stoyan) was computed as K(r) = V / [n(n−1)] × Σ_i_ C(x_i_, r) / ω(x_i_, r), where C(x_i_, r) is the count of other particles within distance r of particle i and ω(x_i_, r) is the fraction of the sphere of radius r centred at x_i_ that lies within the analysis volume, estimated by Monte Carlo sampling of 400 uniformly distributed sphere-surface points. The L function was derived as L(r) = (3K(r) / 4π) − r, such that L = 0 for CSR and L > 0 indicates clustering. The analysis volume was defined as the bounding box of all detected particles, padded laterally in X and Y by the maximum analysis radius (200 nm). The Z dimension was left unpadded because the lamella surfaces form a hard physical boundary on the accessible volume. Statistical significance was assessed against a pointwise 95% Monte Carlo confidence envelope derived from 200 simulations of n particles uniformly distributed within the same bounding box. Bivariate Ripley’s L was computed analogously, counting one particle type within radius r of each particle of the other type, normalised by both particle counts and volume. Analysis distances ranged from 5 to 200 nm in 32 steps; tomograms with fewer than 10 detected particles of a given type were excluded. Analysis was implemented in Python 3 using numpy and scipy.

### PML Body Segmentation and Mesh Generation

PML bodies were manually segmented in Amira using the brush tool (Thermo Fisher Scientific), guided by cryo-CLEM of eYFP-PML-I. Segmentation was performed on IsoNet-deconvolved tomograms (DeconvStrength = 1.5, SnrFalloff = 1.5) to more clearly discern the boundary between the PML body and nucleoplasm. Isosurface volumes of PML bodies were generated and exported as TIFF label files. These segmented volumes were then processed in Python. The PML body label was reduced to a binary mask (body voxels set to one and all other voxels to zero), and the marching cubes algorithm (scikit-image implementation, isovalue threshold = 0.5) was applied to this binary volume to extract triangulated mesh surfaces. The volume data were reoriented by transposing axes (ZYX to XYZ) to match the RELION XYZ coordinate order used in the particle STAR files prior to surface extraction.

The resulting meshes were constructed as trimesh objects containing vertices and face connectivity information, with automatic repair routines applied for non-watertight surfaces including normal fixing, hole filling, and degenerate face removal. For one body (TS_16) the surface was supplied as a pre-exported .obj mesh, already in XYZ order, and loaded directly without the TIFF and marching-cubes steps. Particle coordinates (in unbinned pixels, 2.48 Å/px) were divided by 4 to match the bin-4 mesh coordinate space (9.92 Å/px), and inside/outside classification by ray-casting against the triangulated mesh (trimesh.Trimesh.contains) was performed in this bin-4 pixel space. This classification was applied to the five tomograms with segmented PML bodies (TS_01, TS_03, TS_08, TS_15, TS_16), so the denominators for the reported inside/outside fractions (167 PA28, 194 TRiC, 390 nucleosome and 763 trinucleosome particles) are the counts within these five segmented volumes. Mesh vertices and particle coordinates were scaled to nanometres (0.992 nm/px) only for reporting and rendering. Code implementing this mesh-based visual-proteomics pipeline is available at https://github.com/StephenC07/pml-body-visual-proteomics.

#### Supplementary Note 1: PA28 single-capped proteasome identification by multi-template matching

Seven proteasome-related templates were used for template matching in PyTOM: the 20S core particle (EMDB-4877), single-capped PA28 (EMDB-30824), single-capped PA200 (EMDB-24278), double-capped PA28 (EMDB-24277), double-capped PA200 (EMDB-4860), single-capped 19S (EMDB-4002), and double-capped 19S (EMDB-9512). All templates were pre-aligned in ChimeraX so that the 20S barrel lay along Z and the cap(s) pointed in the same direction, allowing Euler angles from different templates to be directly compared. Template matching across all 16 tomograms yielded 7,795 initial detections: 20S Core (1,468), PA28 (1,659), PA200 (2,335), 19S-Single (402), 19S-Double (1,356), PA28-Double (213), and PA200-Double (362). CC score analysis indicated that 19S templates yielded lower scores than PA28, PA200, and 20S Core detections, while double-capped templates showed comparable scores but substantially fewer detections (Sup. Fig. 11A).

We applied spatial overlap analysis (10 nm tolerance) to group the 7,795 detections into 3,935 unique positions (many detected by more than one template), then used CC-ranked cap assignment to determine the proteasome species at each position, excluding 20S Core detections from the competition. Double-capped templates produced zero winners and 19S templates contributed fewer than 1% of winners, indicating these species were not detected in any of the 16 cryo-tomograms (Sup. Fig. 11C).

To validate cap assignments, we required that each cap detection spatially overlap with a 20S Core particle within 10 nm, and that the cap-Core Y-axis angle was either aligned (< 40°) or flipped (> 140°), consistent with cap binding to either end of the 20S barrel (Sup. Fig. 11B). The orientation filter confirmed the biological validity of the detections: across all inter-template overlapping pairs (n = 8,605), 52.4% fell within the aligned range and 46.5% within the flipped range. Since all templates were aligned with the cap pointing in the same direction, the symmetry of the 20S core barrel means template matching detects it equally in both orientations, producing the observed ∼50/50 split between aligned and flipped cap positions, with only 1.1% falling outside these ranges (Sup. Fig. 11B).

To recover genuine PA28 proteasomes, we required each cap detection to spatially overlap with a 20S Core detection within 10 nm (Phase 3; Sup. Fig. 11C). This two-template filter yielded 875 validated PA28 single-capped proteasome particles and 580 unpaired Core-only detections. STA of the validated PA28 set in RELION 4.0 with iterative 3D classification yielded a final reconstruction of 389 particles at 29 Å resolution (Fig. 2Q–S; Sup. Fig. 12C–F), with clear densities for both the 20S core barrel and the PA28 cap.

